# flowCut — An R package for precise and accurate automated removal of outlier events and flagging of files based on time versus fluorescence analysis

**DOI:** 10.1101/2020.04.23.058545

**Authors:** Justin Meskas, Sherrie Wang, Ryan Brinkman

## Abstract

Technical artifacts that occur during the data acquisition process of cytometry data can result in erroneous data. We showed the presence of these data leads to biased gating analysis. Common technical issues, such as clogging, can cause spurious events and fluorescence intensity shifting. These events should be identified and potentially removed before being passed to the next stage of the gating analysis. flowCut, an R package, automatically detects anomaly events and flags files for flow cytometry experiments. flowCut outperforms existing automated approaches in our evaluation.

flowCut is available as an R package at: https://github.com/jmeskas/flowCut. Test data uploaded to FlowRepository (Repository ID: FR-FCM-ZYPD) along with the manual results.

## 1 Introduction

Even when proper protocols are followed, technical issues such as clogging can result in spurious events in flow cytometry data. Removing those instrumental level problematic events can improve data quality and subsequent downstream analysis. We developed flowCut to address shortcomings and improve performance over the two available automated data cleaning tools (flowAI [1], flowClean [2]). flowClean has a minimum 30,000 cells requirements and, often times, is not able to capturing sufficient abnormal regions, as shown in Figure 1. flowAI can be aggressive, as seen in Figure 1 where flowAI removed a large amount of normal regions in addition to outlier regions. To address the current limitations of the two algorithms, we present a new algorithm, flowCut, as a new state-of-the-art flow cytometry quality control tool. We evaluated flowCut’s performance against that of flowClean and flowAI.

**FIGURE 1:**
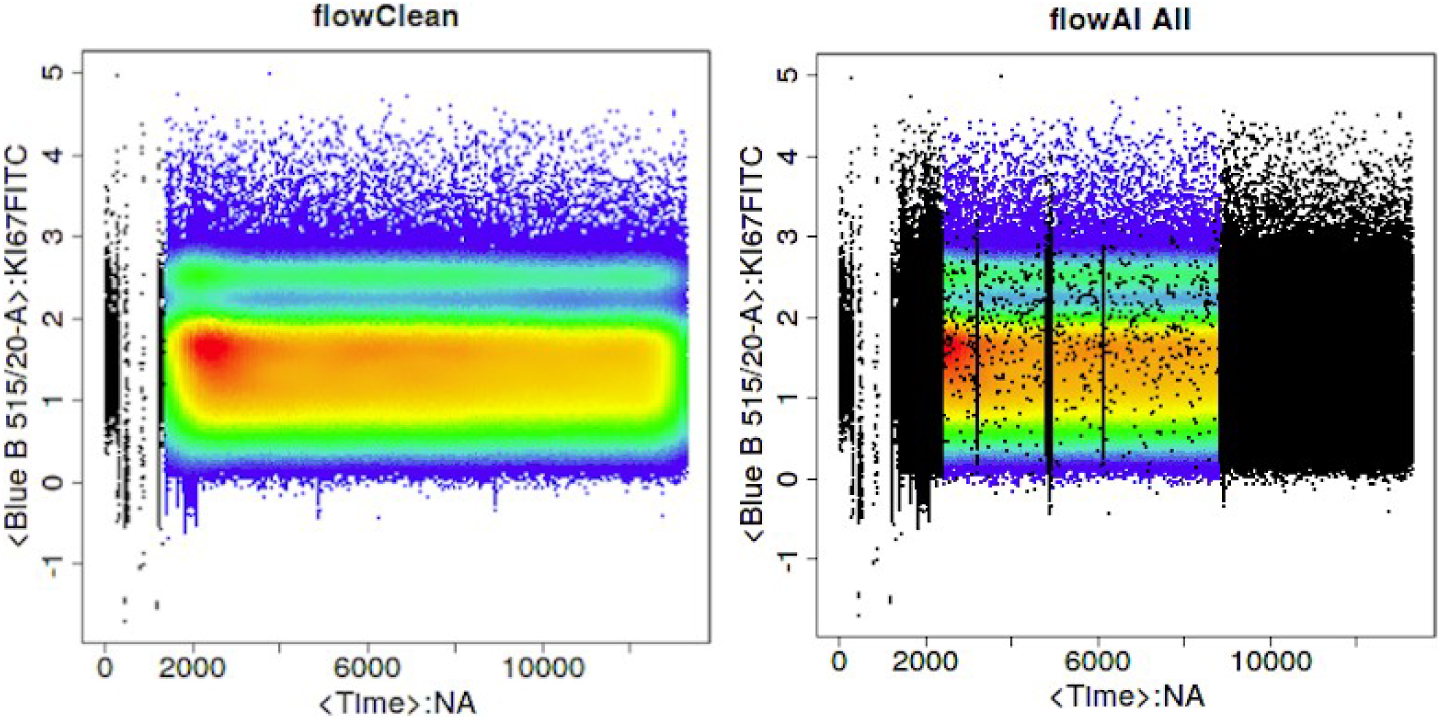
Quality control by flowClean (left) and flowAI (right) on the same file. The colored regions are fluorescence intensity signals. Red is the most dense, followed by yellow, green, and blue. Black is the removed regions.

## 2 Methods

flowCut separates events along the time axis into equally sized segments (parameter defaulted at 500 events) and calculates if any should be removed. This parameter as well as all other numerical user-definable parameters have robust defaults based on extensive testing (described fully in the supplemental materials). flowCut determines if a file needs cleaning based on three tests with user-definable parameters. These three threshold parameters are: the maximum range of the means of each segment across all channels (*MaxOfMeans*), the average range of the means of each segment over all channels (*MeanOfMeans*), and the maximum continuous jump between adjacent segments (*MaxContin*).

Figure 2 (a) shows how the mean drifts a proportion of 0.231 (proportion of the distance between the brown lines, the 2nd and 98th percentiles), where the points of the pink and brown jagged line are the means of the segments. The numbers on top of Figure 2 (a) indicate mean drift before cutting, mean drift after cutting, and the max one step mean change between adjacent segments after cutting. The mean drifts such as the 0.231 are the *Means* when calculating the *MaxOfMeans* and the *MeanOfMeans*. Since *MaxOfMeans* is defaulted at 0.15, the file shown in Figure 2 (a) will undergo cleaning because 0.231 > 0.15.

**FIGURE 2:**
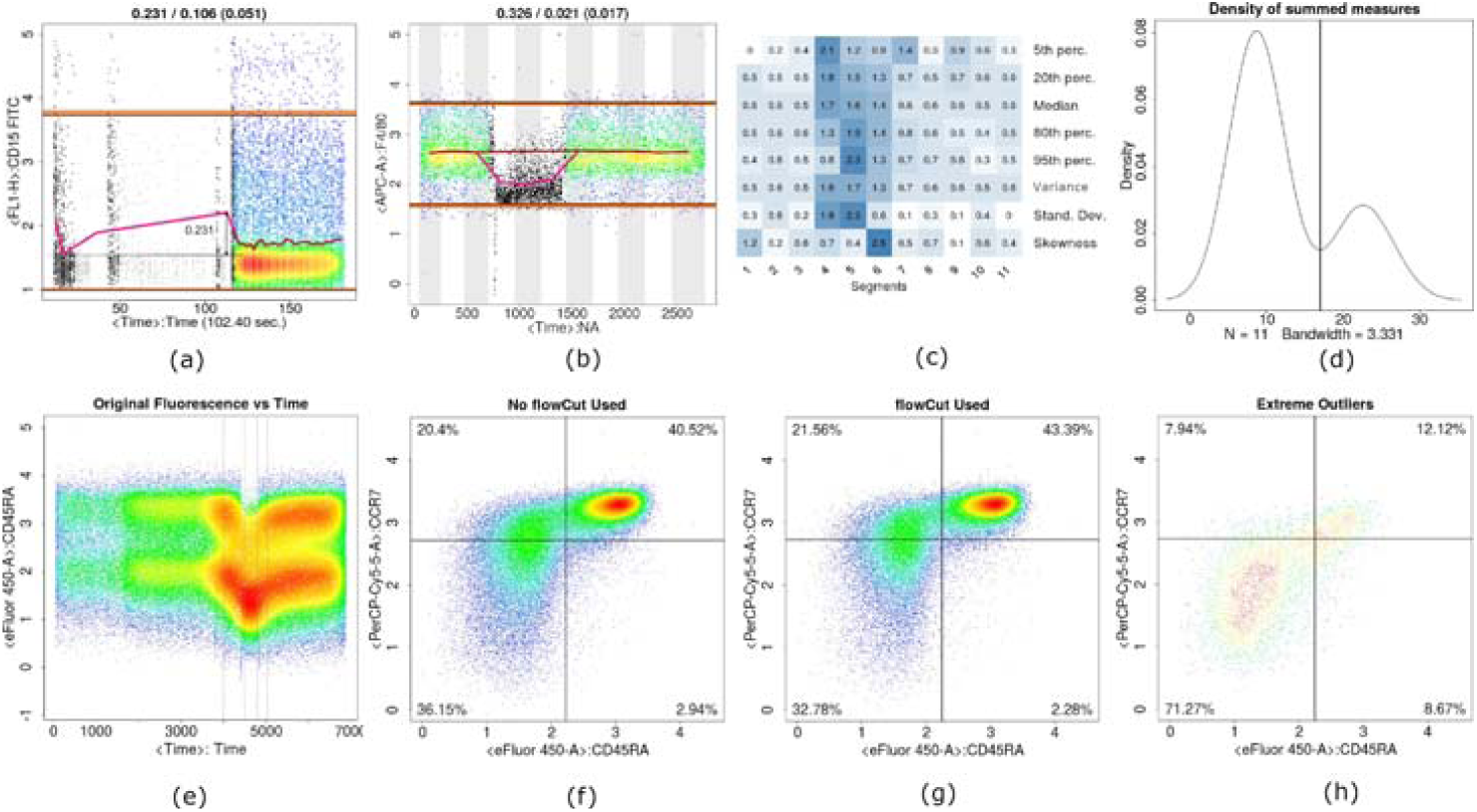
flowCut methodology. (a) flowCut removes low density regions (grey) and statistically different segments (black). (b)-(d) represent the same file. flowCut removes the problematic regions (black in (b)) that have higher statistical values indicated by the darker blue colour in (c) for all eight measures. The 11 columns in (c) correspond to the 11 segments shown in (b) by alternating background colour. We sum over all the measures across all channels to get a statistical score for each segment, and use these 11 values to create a density curve shown in (d). We find a threshold (vertical line in (d)) where the segments with high statistical values, which reside to the right, are removed. (e) shows a new file with fluorescence drifting for CD45RA (Similarly for CCR7 — not shown). (e)-(h) show the difference between (f) not using and (g) using flowCut. (h) shows only the gated events between the middle two grey vertical lines in (e) (the outside grey vertical linesarewhereflowCutremovedevents).

If flowCut deems the file needs cleaning, then eight measures of each segment are calculated (See Figure 2 (c)): the mean, the median, the 5th, 20th, 80th and 95th percentiles, the second moment which accounts for variation and the third moment which accounts for skewness. These measures are absolute Z-scores (absolute because we are only interested in the difference and not the direction). Since the Z-scores are naturally normalized, flowCut can sum the eight measures across all channels and all measures, resulting in each segment being represented by only one statistical number. We utilize the deGate function from flowDensity [3] to find the natural separation point in the distribution to remove outlier segments (See Figure 2 (d)). flowCut then repeats the three tests explained above, and also validates if the file is monotonically increasing in time. If at least one of the four tests are not passed, the file is flagged.

## 3 Tests and Results

We chose 55 exemplary files from FlowRepository [4] to compare the analysis of the three algorithms and a random sampling to manual analysis (see supplementary materials for details). The random sampling was to compare the F1 scores of the algorithms to the F1 score a random cleaning. The percentage of events removed with random sampling was equal to the amount of events removed by manual analysis.

For each of the 55 files, we did blind manual analysis that visually identified the problematic regions in the fluorescence vs time domain in every channel after compensation, transformation and margins removal. Problematic regions include spikes, fluorescence drifting, discontinuity or low density.

### 3.1 Comparing Algorithms with Manual Gating — File Based Evaluation

We then subdivided the 55 files into three categories based on confidence in the manual analysis. The first category includes 17 files that have removal regions with clearly defined boundaries such as discontinuity, low density regions and large spikes. The second category has 35 files that have fuzzy boundaries for removal regions. Examples include small spikes and cutoff in fluorescence drifting regions. This category also includes files that have overlapping problematic regions detected by at least two algorithms but not by manual analysis. The third category has 3 files which manual analysis is arbitrary. The files look abnormal but the regions for removal are not clear (see supplementary material). Category three files contain no region of truth and are removed from being used as standards for evaluation. For the remaining 52 files, we calculated F1 scores for two categories for each algorithm.

The F1 score is 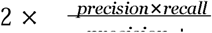, where precision is defined as the proportion of each algorithm’s intersection with manually removed events in total manually removed events; recall is defined as the proportion of each algorithm’s intersection with manually removed events in all events removed by each individual algorithm. The results of the manual gating were treated as the standard for computing the F1 scores. Manual analysis took approximately 5 minutes per file. The results of category one files and category two files are shown in Table 1 and Table 2, respectively.

**TABLE 1:**
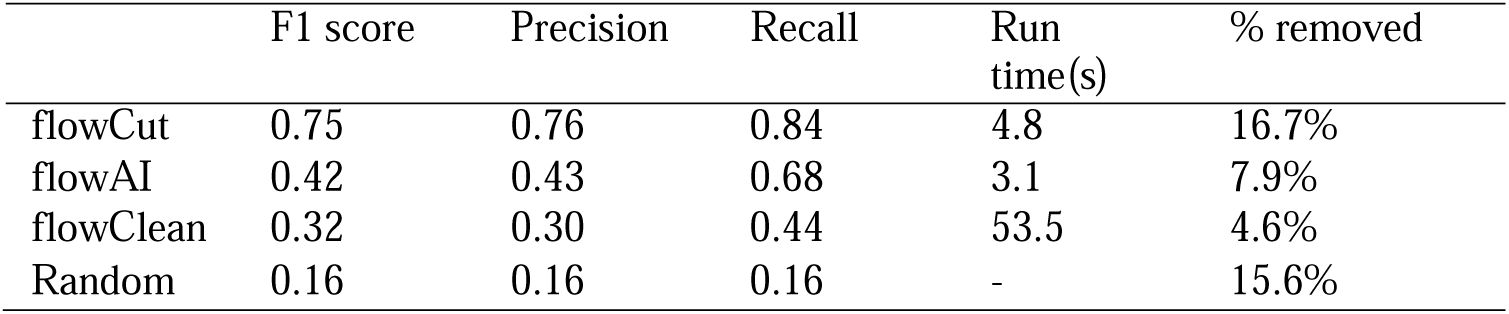
F1 scores, precision, recall, run times and percentage removed of each algorithm when comparing to in-house manual analysis for 17 **category one** FlowRepository exemplary files. Computed on an Intel Xeon E5-2630 CPU with 128 GB RAM.

|  | F1 score | Precision | Recall | Run<br>time(s) | % removed |
| --- | --- | --- | --- | --- | --- |
| flowCut | 0.75 | 0.76 | 0.84 | 4.8 | 16.7% |
| flowAI | 0.42 | 0.43 | 0.68 | 3.1 | 7.9% |
| flowClean | 0.32 | 0.30 | 0.44 | 53.5 | 4.6% |
| Random | 0.16 | 0.16 | 0.16 | - | 15.6% |

**TABLE 2:**
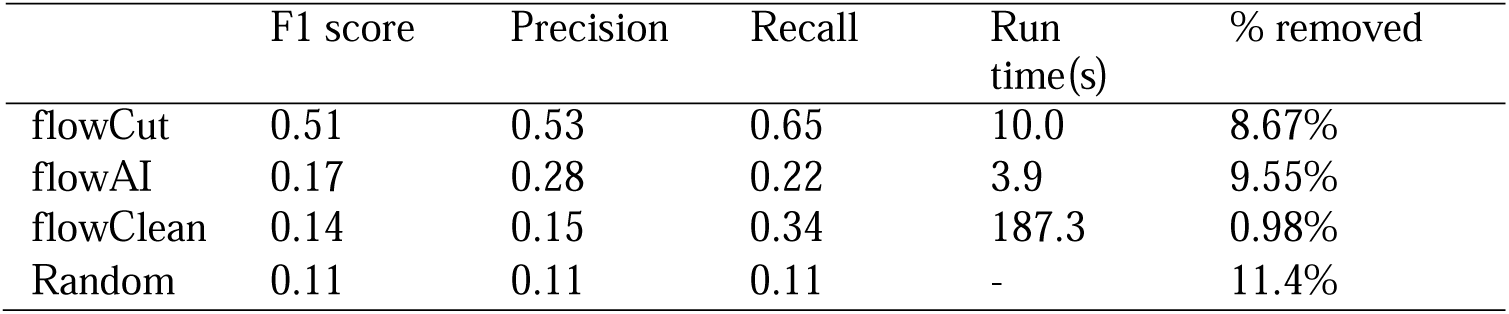
F1 scores, precision, recall, run times and percentage removed of each algorithm when comparing to in-house manual analysis for 35 **category two** FlowRepository exemplary files. Computed on an Intel Xeon E5-2630 CPU with 128 GB RAM.

|  | F1 score | Precision | Recall | Run<br>time(s) | % removed |
| --- | --- | --- | --- | --- | --- |
| flowCut | 0.51 | 0.53 | 0.65 | 10.0 | 8.67% |
| flowAI | 0.17 | 0.28 | 0.22 | 3.9 | 9.55% |
| flowClean | 0.14 | 0.15 | 0.34 | 187.3 | 0.98% |
| Random | 0.11 | 0.11 | 0.11 | - | 11.4% |

### 3.2 Comparing Algorithms with Manual Gating - Problem Based Evaluation

We categorize all problematic regions into four types: shift, spike, low density and discontinuity. Each type has its unique characteristics. For example, mean shift is characterized by a gradual transition of means into a different position, spike is characterized by an abrupt jump between adjacent segment means, the size of which is small. Low density is having very few cells in any periods of time. Discontinuity is an abrupt change of means between two or more large sections of data. We evaluate algorithms’ performance on dealing with these four types of problems by calculating F1 scores for all regions of each type. We also calculate F1 scores on all clean regions to capture false positives and false negatives. The mean F1 scores for each problematic type with their normalized percentage of events in a file are shown in Table 3. The percentage of events is calculated as the sum of all regions of that type identified by manual analysis divided by the total number of files (52). The sum of the products of mean F1 scores of each type and their proportions in a file is the weighted F1. These weighted F1 scores are flowCut 0.93, flowAI 0.86 and flowClean 0.83 (Table 3).

**TABLE 3:** Mean F1 scores of four types of problematic regions, mean percentage of events by manual analysis and the weighted F1 scores for all three algorithms

|  | % of events | flowCut | flowAI | flowClean |
| --- | --- | --- | --- | --- |
| Shift | 7.23% | 0.67 | 0.40 | 0.20 |
| Spike | 2.60% | 0.63 | 0.33 | 0.02 |
| Low Density | 1.72% | 0.67 | 0.28 | 0.20 |
| Discontinuity | 0.98% | 0.93 | 0.32 | 0.00 |
| Clean | 87.47% | 0.96 | 0.93 | 0.93 |
| Weighted F1 | - | 0.93 | 0.86 | 0.83 |

### 3.3 flowCut’s Robustness

While searching for the 55 exemplary files, we had crashes or unrecoverable errors on 0%, 6% and 11% of the files for flowCut, flowAI and flowClean, respectively. In addition, we ran flowCut on all 117,115 FlowRepository’s public files and there were 0 files with crashing or unrecoverable errors (October 2019).

### 3.4 Biased gating populations

We reproduced the gating of CD45RA-CCR7+ TCM CD8 T cells with and without using flowCut using a publicly available dataset [5] (FlowRepository ID: FR-FCM-ZYBU). The data that has undergone proper data quality control shows an increase from 20.4% to 21.56% of CD45RA-CCR7+ (Figure 2 (e)-(h)). The two outer grey vertical lines in Figure 2 (e) mark the location of where flowCut started and ended it’s cleaning. The two central vertical grey lines are set manually to capture the extreme outliers to show emphasis in Figure 2 (h).

## 4 Discussion

Data cleaned by flowCut improves the downstream gating. flowCut allows users to check the quality of the cleaning and to adjust stringency of the algorithm if needed. Compared to existing methods, flowCut identifies outlier events more accurately and does not fail to process any file.

## Supporting information

flowCut vignette

paper_code

## Acknowledgements

Special thanks to Sibyl Drissler and Jessica Tuengel for their testing, feedback and criticism.

Research reported in this publication was supported in part by NSERC, NIGMS [R01 GM118417-01A1], an NIH Infrastructure and Opportunity Fund Award linked to Human Immunology Project Consortium Award Number [U19AI118610] (IOF Pilot), and the Precision Vaccines Program.

The content here only represents the views of the authors and does not necessarily reflect the official views of the NIH.

## List of Figures

1. Quality control by flowClean (left) and flowAI (right) on the same file. The colored regions are fluorescence intensity signals. Red is the most dense, followed by yellow, green, and blue. Black is the removed regions. 10
2. flowCut methodology. (a) flowCut removes low density regions (grey) and statistically different segments (black). (b)-(d) represent the same file. flowCut removes the problematic regions (black in (b)) that have higher statistical values indicated by the darker blue colour in (c) for all eight measures. The 11 columns in (c) correspond to the 11 segments shown in (b) by alternating background colour. We sum over all the measures across all channels to get a statistical score for each segment, and use these 11 values to create a density curve shown in (d). We find a threshold (vertical line in (d)) where the segments with high statistical values, which reside to the right, are removed. (e) shows a new file with fluorescence drifting for CD45RA (Similarly for CCR7 — not shown). (e)-(h) show the difference between (f) not using and (g) using flowCut. (h) shows only the gated events between the middle two grey vertical lines in (e) (the outside grey vertical lines are where flowCut removed events) 11

## List of Tables

1. F1 scores, precision, recall, run times and percentage removed of each algorithm when comparing to in-house manual analysis for 17 **category one** FlowRepository exemplary files. Computed on an Intel Xeon E5-2630 CPU with 128 GB RAM.13
2. F1 scores, precision, recall, run times and percentage removed of each algorithm when comparing to in-house manual analysis for 35 **category two** FlowRepository exemplary files. Computed on an Intel Xeon E5-2630 CPU with 128 GB RAM.14
3. Mean F1 scores of four types of problematic regions, mean percentage of events by manual analysis and the weighted F1 scores for all three algorithms 15

## References

[1] flowAI: automatic and interactive anomaly discerning tools for flow cytometry data. Bioinformatics. 2016 08;32(16):2473–2480.

[2] flowClean: Automated identification and removal of fluorescence anomalies in flow cytometry data. Cytometry A. 2016 05;89(5):461–471.

[3] Malek M, Taghiyar MJ, Chong L, Finak G, Gottardo R, Brinkman RR. flowDensity: reproducing manual gating of flow cytometry data by automated density-based cell population identification. Bioinformatics. 2015;31(4):606–607.

[4] FlowRepository: a resource of annotated flow cytometry datasets associated with peer-reviewed publications. Cytometry A. 2012 Sep;81(9):727–731.

[5] An Integrated Workflow To Assess Technical and Biological Variability of Cell Population Frequencies in Human Peripheral Blood by Flow Cytometry. J Immunol. 2017 02;198(4):1748–1758.

