## Supplementary material for "flowCut — An R package for precise and accurate automated removal of outlier events and flagging of files based on time versus fluorescence analysis"

### flowCut — Supplementary Material

**Running head:** flowCut — Supplementary Material

[*]

[†]

[‡]

### 1 Parameters

An in-depth discussion of all main parameters is provided in the separate vignette.

### 2 Outliers example

The paper includes examples of three types of outlier events: low density, fluorescence drifts and fluorescence discontinuity. Here we show the fourth type of outlier: spikes. Supplementary figure 1 shows multiple spiking at the beginning and one spike near the middle of the data acquisition. The sub-figure on the left is the uncleaned data, while the sub-figure on the right shows the removed sections in black.

[Supplementary

figure 1 about here.]

### 3 deGate( ) Steps

- flowCut first finds all the peaks (p = 1, 2, ...n) in the density distribution.
- If p >= 2, flowCut calculates the height difference from the valley(s) to its smaller adjacent peak. If it is less than one percent of the height difference between the valley(s) and the higher adjacent peak, then flowCut ignores the lower peak.
- If there are still two or more peaks left after the previous step and the proportion of the population to be removed is less than or equal to the user specified amount of *MaxPercCut* (a parameter that sets an upper bound of which percentage of events can be removed), flowCut will remove the significantly different population.
- If p = 1, it uses the deGate function to find a natural point (the fastest derivative

change) along the upstream of the density distribution to remove significantly different segments.

### 4 55 files

For comparing the three methods, we randomly downloaded 1071 FCS files from FlowRepository.
We discarded 83 files that were corrupted or had issues with compensation or transformation. There were 145 files that resulted in an unrecoverable error from at least one algorithm. flowClean uniquely crashed for 80 files, while flowAI uniquely crashed 31 files and flowCut 0 files. flowClean and flowAI both also crashed an additional 34 files. We also discarded 788 files that had only one or no algorithm that cleaned some events. However, we made an exception for five files that had only one algorithm cleaning and one file that had zero algorithms cleaning. These six files were noticeably requiring cleaning, and hence we decided to keep them.

The resulting 55 files were used to compare the analysis of the three algorithms to manual analysis. These 55 files were uploaded to FlowRepository, and can be found with Repository ID of FR-FCM-ZYPD. (https://flowrepository.org/id/FR-FCM-ZYPD)

### 5 Algorithm versions

Comparisons were done with flowCut 0.99.26, flowAI 1.18.1 and flowClean 1.26.0 all on default settings except in flowAI's function, *flow_auto_qc( )* we used *remove_form='FS_FM'* since we did not want to exclude events due to changes in flow rate. From our knowledge flowrate plays little to no affect on the fluorescence readings of the events (with the exception of very high flowrates). Also, there are many times when flow cytometer users adjust the flowrate on the fly for various reasons.

Other packages used and their versions are as follows: flowCore 1.51.7, flowDensity

1.20.0, doMC 1.3.6, foreach 1.4.8, Cairo 1.5-10, FlowRepositoryR 1.14.1, R.utils 2.9.2 and stringr 1.4.0.

### 6 Code

The code used to calculate the F1 scores is available as supplementary materials. The code is also available on FlowRepository with Repository ID of FR-FCM-ZYPD. (https://flowrepository.org/id/FR-FCM-ZYPD). This code will automatically download the data from FlowRepository and re-create the results.

### 7 FlowRepository experiments of the 55 files

The following list shows which FlowRepository experiment the 55 exemplary files came from. They are divided into three categories. (These files have been re-uploaded to FlowRepository with Repository ID of FR-FCM-ZYPD).

#### 7.1 Category one files:

- FR-FCM-ZZ5V/2 dias_Infectado 5.fcs
- FR-FCM-ZZ99/003.fcs
- FR-FCM-ZZ4L/9407_2_1_NKR.fcs
- FR-FCM-ZZEX/-_BDL aLFA-1.fcs copy
- FR-FCM-ZY7H/binding assay_dilution 6_007.fcs
- FR-FCM-ZZ8S/GDC-0941-CQ_D10_D10.fcs
- FR-FCM-ZZKH/IL2-20_pepstim#1 1501.fcs-(1)
- FR-FCM-ZZ7E/Macrophages + Leishmania + oATP.fcss
- FR-FCM-ZZ7E/Macrophages + Leismania.fcs
- FR-FCM-ZZ7E/Macrophages + oATP.fcs
- FR-FCM-ZZUL/PBL_7wk M5.fcs
- FR-FCM-ZZVB/Specimen_001_B6 LSK.fcs

- FR-FCM-ZZ8L/TH-004_TH-004 mg.fcs

- FR-FCM-ZZF5/TS05 121 P6.fcs

- FR-FCM-ZZFE/TS14 351 P6.fcs

- FR-FCM-ZZFS/TS27 937 P4.fcs

- FR-FCM-ZZ8F/Unstained SPL_C3_C03.fcs

#### 7.2 Category two files:

- FR-FCM-ZZD7/100_111 SEB.fcs

- FR-FCM-ZZD7/100_111 vehicle.fcs

- FR-FCM-ZZD7/125_114 Pre_6b.fcs

- FR-FCM-ZZ2T/13523 17012011_NS_F01.fcs

- FR-FCM-ZZ88/15_24.fcs
- FR-FCM-ZZ45/2151_074_BA2012-02-28_020.fcs
- FR-FCM-ZZ45/2151_074_BA2012-02-28_021.fcs
- FR-FCM-ZY6K/2nd group-high amylose maize_6_006.fcs
- FR-FCM-ZZVD/50uM ALLN-Asy48h.fcs
- FR-FCM-ZYBN/6A_liver_3d_WT.fcs
- FR-FCM-ZZ88/7c_MA+.fcs
- FR-FCM-ZZ4L/9399_2_1_NKR.fcs
- FR-FCM-ZZ4L/9399_2_4_NKR.fcs
- FR-FCM-ZZ5D/9606_3_9_NKR.fcs
- FR-FCM-ZZ9P/ah20130417 dilution_Tube_025 surface_Sup 2.fcs
- FR-FCM-ZZ72/B3 C.reinhardtii H2O2 5mM_ stained 2.5 microM C11-BODIPY581_591 .fcs

- FR-FCM-ZZ72/D33 C.reinhardtii preloaded 2.5 microM C11-BODIPY581_591+H2O2 10 mM t=25.fcs
- FR-FCM-ZZ7N/Fig4_Algae_chlorination.fcs
- FR-FCM-ZZ7E/Macrophages.fcs
- FR-FCM-ZZ8Z/Paciente AH18_Isotipos.fcs
- FR-FCM-ZZMJ/PBMC_HuK-CD200-14.fcs
- FR-FCM-ZZYS/PBMC_Tphe_PROP10813_P4_35_T48B_E08.fcs
- FR-FCM-ZZK3/Phytoplankton_shallow sample.fcs
- FR-FCM-ZZGS/Tphe09943-006-00_F7_R.fcs
- FR-FCM-ZZEV/TS01 14 P7.fcs
- FR-FCM-ZZEV/TS01 25 P5.fcs

- FR-FCM-ZZF4/TS04 92 P6.fcs
- FR-FCM-ZZF6/TS06 137 P3.fcs
- FR-FCM-ZZF6/TS06 144 P5.fcs
- FR-FCM-ZZFG/TS16 406 P7.fcs
- FR-FCM-ZZFL/TS21 802 P2.fcs
- FR-FCM-ZZFL/TS21 813 P5.fcs
- FR-FCM-ZZFS/TS27 952 P2.fcs
- FR-FCM-ZZKH/UR414 24_Direct ex vivo 1502.fcs
- FR-FCM-ZZ8L/VD_TH-003.fcs

#### 7.3 Category three files

- FR-FCM-ZZVU/2014-04-24_T18_T_cell_panel_T_cells_CD45RO-164-new_CD45tight.fcs
- FR-FCM-ZZZA/CD34+CD38- and CD34+CD243+ populations in human Term Cord Blood.fcs
- FR-FCM-ZZJ7/GvHD8.fcs

Note: The 12 file names with the following syntax "TS_______P_", where the " _ " is a number, have had their file names renamed since they used to contain unrecognized characters. The updated file names are easily matched with the old file names inside their respective experiments.

Category three files are shown in Supplementary figure 2. They are categorized as such because the manual analysis can be arbitrary. Supplementary figure 2 (a) contains three sections of mean drifts. It is not clear which section(s) should be best removed. Supplementary figure 2 (b) has two equally proportioned population sections with different means. It is not clear which section one should remove. Supplementary figure 2 (c) also contains similar ambiguity. Manual analysis on these files are not good standards for evaluating algorithms. Therefore, all evaluations done in this paper are based on category one and two files only.

[Supplementary

figure 2 about here.]

### 8 Manual gating procedure

We plot every channel versus time for each file and visually identify problematic regions. Each region to be removed is defined by two boundaries; the beginning and the end. These boundaries have been listed in Manual_Gating_52.csv, which can be found in the FlowRepository experiment FR-FCM-ZYPD (https://flowrepository.org/id/FR-FCM-ZY PD). Each row in this csv file represents one of the 52 files used (3 files are excluded due to uncertainty in the manual analysis). Each row has manual gates that are comprised of a series of an even amount of boundaries that define the regions for cutting. For example, if there are four numbers, the first and second numbers are the beginning and end of the first region removed. And the third and fourth values are the beginning and end of the second region removed. We subsequently categorize all regions in manual analysis into four types. The categorized regions are saved in Manual_Gates_byProblemtype.csv.

### List of Figures

1 Example of spikes. The left plot is uncleaned and the right plot shows flowCut's removed events in black....................................................................9

2 The worst channel of each of the category three file is shown.........................10

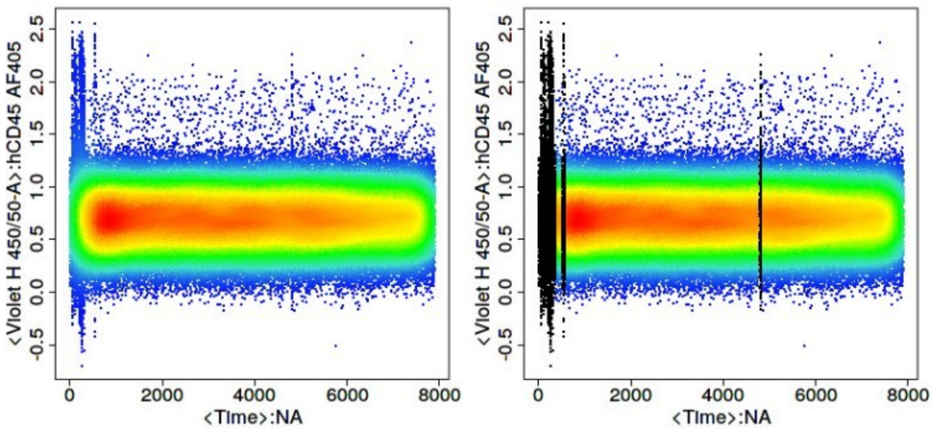


SUPPLEMENTARY FIGURE 1: Example of spikes. The left plot is uncleaned and the right plot shows flow- Cut’s removed events in black.

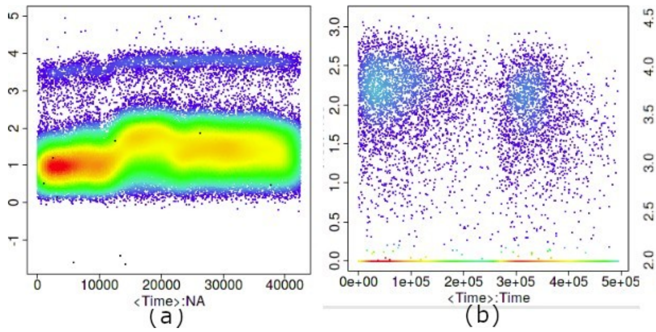


SUPPLEMENTARY FIGURE 2: The worst channel of each of the category three file is shown.
