## Supplementary material for "flowCut — An R package for precise and accurate automated removal of outlier events and flagging of files based on time versus fluorescence analysis": flowCut vignette

### License

Artistic-2.0

### Package from GitHub

***flowCut*** can be downloaded from "https://github.com/jmeskas/flowCut" or cloned from "https://github.com/jmeskas/flowCut.git". For more information on installation guidelines, see the GitHub, Bioconductor and CRAN websites. Once installed, to load the library, type the following into R:

```
library("flowCut")
```

### Running *flowCut*

#### Introduction

***flowCut*** automatically removes outlier events in flow cytometry data files due to abnormal flow behaviours resulting from clogs and other common technical problems. Our approach is based on identifying both regions of low density and segments (default size of 500 events) that are significantly different from the rest.

Eight measures of each segment (mean, median, 5th, 20th, 80th and 95th percentile, second moment (variation) and third moment (skewness)) are calculated. The density of the summation, over both the 8 measures of each segment and all cleaning channels, gives a density of summed measures distribution. This distribution, accompanied with two parameters (*MaxValleyHgt* and *MaxPercCut*), will determine which segments are significantly different from the other segments. The segments that are significantly different will have all of their events removed. We also flag files if they display any of the following: 1) not monotonically increasing in time, 2) sudden changes in fluorescence, 3) large gradual change of fluorescence in all channels, or 4) very large gradual change of fluorescence in one channel. (During this vignette four digit codes of only Ts and Fs will be seen, ex. TFTF. These codes are a pass/flag of the 4 flagging tests).

#### Data

The ***flowCut*** dataset contains flowFrames borrowed from the *flowCore's* GvHD dataset and from FlowRepository, an open source cytometry data repository. The first two flowFrames are GvHD flowFrame objects 's9a06' and 's5a07'. The third flowFrame is from FlowRespository's experiment FR-FCM-ZZ7E with file name 'Macrophages + oATP.fcs'. The fourth is from FlowRespository's experiment FR-FCM-ZZVB with file name 'Specimen_001_B6 LSK.fcs'. We pre-loaded the compensated and transformed data into the ***flowCut*** library. To load all the data used in the examples of this vignette, type:

```
data(flowCutData)
```

#### Simple Example

We run ***flowCut*** with default parameters with the exception of *FileID* that gives a name to the figure and *Plot* that is changed to 'All' so we can see the results even if the file is perfectly cleaned. The default for *Plot* is 'Flagged Only' which would only plot the flagged files. Throughout this vignette all the examples will produce a figure in a folder called flowCut in your current directory. As an alternative *PrintToConsole* can be changed to 'TRUE' which allows the image to be printed in R as opposed to being saved as a file. If you would prefer this alternative, simply add *PrintToConsole=TRUE* to all ***flowCut*** commands.

```r
res_flowCut <- flowCut(flowCutData[[1]], FileID = 1, Plot = "All")
```

A list containing four elements will be returned. The first element, *$frame*, is the flowFrame file returned after ***flowCut*** has cleaned the flowFrame. The second element, *$ind*, is a vector containing indices of segments being removed, in relation to the original flowFrame. The third element, *$data*, is a table containing computational information that users can access. The fourth element, *$worstChan*, is the index of the channel with the most drift over time before cleaning. The *$data* element is shown here as an example:

```r
res_flowCut$data
```

```
##                                                        [,1]
## Is it monotonically increasing in time                 "T"
## Largest continuous jump                                "0.051"
## Continuous - Pass                                      "T"
## Mean of % of range of means divided by range of data "0.075"
## Mean of % - Pass                                       "T"
## Max of % of range of means divided by range of data  "0.114"
## Max of % - Pass                                        "T"
## Has a low density section been removed                 "T"
## % of low density removed                               "5.32"
## How many segments have been removed                    "4"
## % of events removed from segments removed              "12.04"
## Worst channel                                          "FL1-H"
## % of events removed                                    "17.36"
## FileID                                                 "1"
## Type of Gating                                         "MaxPercCut"
## Was the file run twice                                 "No"
## Has the file passed                                    "T"
```

Having the *Plot* parameter set to 'All' above created a plot in the flowCut folder under your current working directory, which is the same as Figure 1. The first five sub-plots in Figure 1 are the channels that undergo fluorescence analysis from ***flowCut***. The segments that are removed because they are significantly different are coloured black. The removed low density sections lower than *LowDensityRemoval*, defaulted at 0.1 (10%), are coloured grey. The low density sections are calculated by first creating a density distribution. Then the parts of the distribution curve that are less than 10% of the max value are removed. This is independent of the rest of the algorithm and is done first and has no association to the binning of the segments. In other words, any number of events can be removed and not just an integer amount of the segment size. On the other hand, events removed from being part of a significantly different segment are removed altogether as a segment and no one single event could be removed.

The top and bottom horizontal dark brown lines correspond to the 98th and the 2nd percentile of the data after cleaning. The 98th and 2nd percentile lines for the full data before cleaning are coloured in light brown, but are most likely not seen. Most of the time these two sets of lines are indistinguishable on the plot because there is no significant change in percentiles before and after cleaning. We found the top and bottom 2% of the events are spread out too much and do not serve as a nice max and min value of the file when comparing the range of the means to the range of the file.

Connecting all the means of each segment gives the brown line in the middle. Sometimes you can see that the brown line has some pink parts. The pink parts are the mean of each segment before cleaning. With the difference between pink and brown, the user can see which segments are removed, and perhaps understand why as well. Of course this only shows the mean measure of the 8 measures used to judge if a segment is removed or not, so information on the other 7 measures is not shown.

The numbers on top of each of the eight plots (in the form of A / B ( C ) ) indicate mean drift before cutting (A), mean drift after cutting (B) and the max one step change after cutting (C). The mean drift is

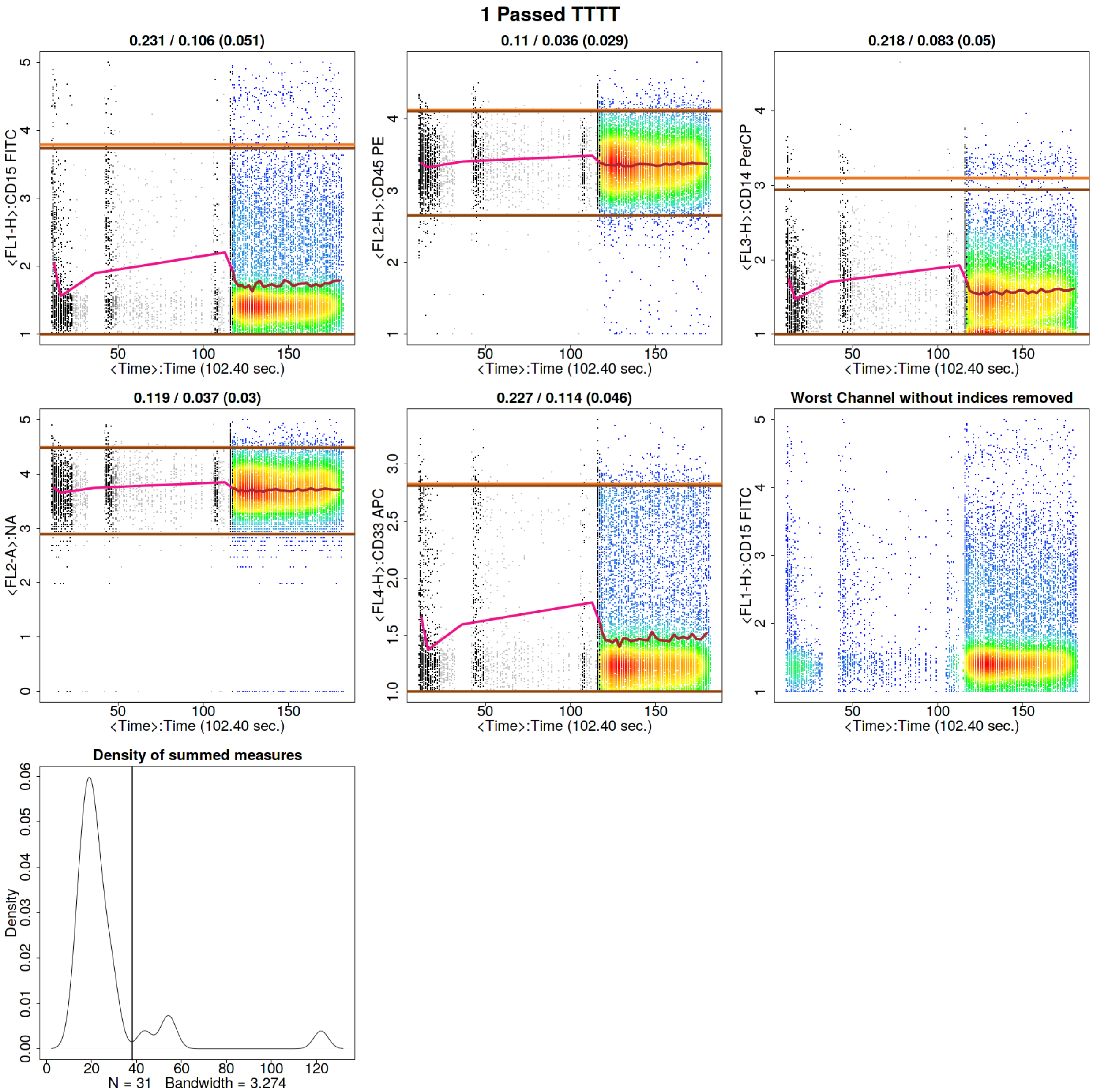


Figure 1: Running ***flowCut*** with the default values.

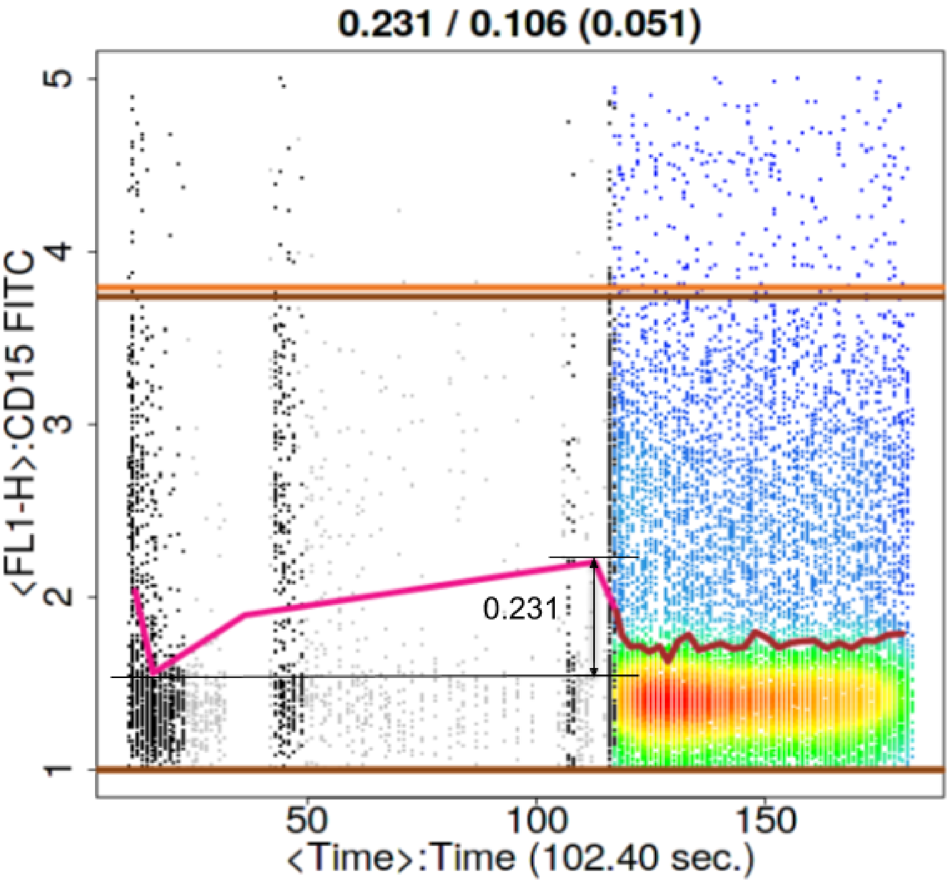


Figure 2: How MaxOfMeans is calculated.

calculated as the difference of the maximum mean and the minimum mean divided by the 2-98 percentile difference of the data. This would catch any gradual changes or fluctuations in the file. If the mean drift becomes significant, the file will be attempted to be cleaned (pre-cleaning) or flagged (post-cleaning). The parameters *MeanOfMeans* and *MaxOfMeans* control the strictness of these flags, and we recommend not to change these numbers unless a greater or weaker strictness is desired. The *MeanOfMeans* parameter is the threshold for the average of the range of means of all channels, where *MaxOfMeans* is the threshold just for the channel with the highest range of means.

Figure 2 illustrates what the range of the means of the segments on the CD33 channel from Figure 1. We will explain the ideas in the previous paragraph with this picture in mind. The range of 0.231 from Figure 2 is the difference between the maximum and the minimum mean of the segments in one channel. If this ranged value for the largest channel is larger than *MaxOfMeans* the file will undergo cleaning. If the average of this ranged value for all markers is larger than *MeanOfMeans* the file will undergo cleaning. In other words, *MeanOfMeans* and *MaxOfMeans* essentially set a restriction for a tolerable mean drift.

The value in the bracket (C) is associated to the parameter *MaxContin*, which indicates the changes of the means between adjacent segments in each channel. This parameter would catch abrupt mean changes such as spikes. The default is 0.1 and we recommend not to change this value. Saying it again in a different way, if adjacent segments have differences in their means that exceed *MaxContin*, then the file will undergo cleaning.

If, after cleaning, the file still has at least one larger value for one of the threshold parameters (*MaxOfMeans*, *MeanOfMeans*, *MaxContin*), then the file will be flagged. Hence, these three parameters have the ability to judge first if the file needs to be cleaned, and second if it is cleaned sufficiently enough to not get flagged afterwards.

The 8 statistical measures ***flowCut*** calculates from each segment are summed up (and over all cleaning channels) to obtain a single statistical number for each segment. If we plot the density of these summed measures for every segment, then we obtain the last plot in Figure 1. The *deGate* function in *flowDensity* gives us the cutoff line on the density curve (This is discussed in detail in the next section). Any segment with a summed value larger than the cutoff line will be removed.

The title of the image will display if the file passed or it was flagged (This information is also visible in *$data*). In Figure 1 a '1 Passed TTTT' is displayed. The '1' is for the file's ID, 'Passed' is stating the file passed and 'TTTT' is showing that it has passed all flagging tests (please see Introduction for the four flagging tests).

#### Using deGate from flowDensity to find the gating threshold

Generally, the deGate function returns a gating value of a 1D density profile. This was originally designed to separate cell populations. However, it could be utilized in our outlier detection methodology. We manipulated the deGate function so that it will always return a gating line to the right of the majority of events (or the highest peak) because we are only interested in removing the cells that are most different. Since the density of summed measures distribution is of Z-scores, naturally the significantly different segments lie to the right of the distribution.

The size of the second biggest peak will play a role in the gating process that we use. If the second biggest peak is less than 10% of the height of the biggest peak, then the file will be treated as a uni-modal distribution. This is achieved by setting the *tinypeak.removal* parameter in deGate to 0.1. For these uni-modal distributions, we want to calculate a gating threshold to the right of the highest peak that naturally separates the segments in the density of summed measures distribution. Therefore, we want to find a valley that is close to zero in the density distribution. To do this, we first allow deGate to search for valleys that have at least 0.1% (*tinypeak.removal* = 0.001) of the height of the highest peak, and then make sure they have a very short valley. Allowing for a very short valley or a tall one can be controlled by adjusting the *MaxValleyHgt* parameter. We find setting it to 0.1 forces the algorithm to always find the best natural separation in the density of summed measures distribution. Details on how changing this parameter will affect results is discussed in the next section. There are times when the second biggest peak is larger than 10% of the height of the highest peak. These cases we label as bi-modal, and calculating the gating line between the two peaks is quite straight forward using deGate.

There are two additional cases that would stop the algorithm from cleaning a file that seems to have a clearly separable distribution. The first is that ***flowCut*** has a preliminary checking step. If the file passes all the flagging tests (values less than *MaxContin*, *MeanOfMeans* and *MaxOfMeans*), then it is deemed to already be clean, and therefore does not need to be cleaned. In this case, the density of summed measures distribution could be clearly bi-modal, however, the file is clean enough. If you still want the file cleaned you can decrease one of *MaxContin*, *MeanOfMeans* or *MaxOfMeans* low enough to get the file marked as needing cleaning. The second case preventing ***flowCut*** from removing the amount of events you would expect to be removed is the user defined parameter *MaxPercCut*. This essentially tells the algorithm what the maximum percentage of events the user is willing to cut off. If the percentage of events to the right of the gating threshold exceeds *MaxPercCut*, then ***flowCut*** will not use this gating threshold (an exception of this exception is discussed in the Section called 'Changing the Parameter *MaxPercCut*'). ***flowCut*** will then attempt to find a gating threshold that would remove less than the *MaxPercCut* amount.

#### Changing the Parameter *MaxValleyHgt*

This example will show what *MaxValleyHgt* does and how changing the value will affect the segment deletion analysis.

The *MaxValleyHgt* parameter plays a role with calculating the threshold on the density of summed measures distribution. *MaxValleyHgt* defines the upper limit of the ratio of the height of the intersection point where the cutoff line meets the density plot and the height of the maximum peak. Setting the number higher can potentially cut off more events, whereas lowering the number will potentially reduce it.

For example, if we set *MaxValleyHgt* to 0.01 we have:

```
res_flowCut <- flowCut(flowCutData[[2]], FileID=2, Plot ="All", MaxValleyHgt = 0.01)
```

The file is flagged because the statistically different regions are not completely removed, shown in Figure 3. There are three small bumps in the 'Density of summed measures' sub-plot, which each represent one segment. Also note, that the threshold is set at around 45. If we change the *MaxValleyHgt* back to its default value 0.1, we get Figure 4. Here the file has now passed and is cleaned properly. note that the threshold is now around 33, and hence all three bumps/segments are removed:

```
res_flowCut <- flowCut(flowCutData[[2]], FileID=3, Plot ="All", MaxValleyHgt = 0.1)
```

#### Changing the Parameter *MaxPercCut*

Sometimes we get files that show a distinctive separation of good and undesired segments, as shown in the 'Density of summed measures' sub-plot in Figure 5. We first show *MaxPercCut* changed to 0.15 as a demonstration:

```
res_flowCut <- flowCut(flowCutData[[3]], FileID=4, Plot ="All", MaxPercCut = 0.15)
```

*MaxPercCut* defines the maximum percentage of events allowed to be removed. The default value is 0.3, which means ***flowCut*** can remove 30 percent of the total events. However, with setting it at 0.15 we did not let ***flowCut*** remove any segments since ***flowCut*** needs to remove more than 15%. If we change it back to the default of 0.3 we have a result shown in Figure 6:

```
res_flowCut <- flowCut(flowCutData[[3]], FileID=5, Plot ="All", MaxPercCut = 0.3)
```

With *MaxPercCut* set at 0.3, a large section is removed. This large section is actually much bigger than 30%.

```
res_flowCut$data["% of events removed",]
```

```
## % of events removed
##              "42.27"
```

***flowCut*** could potentially remove more depending on the nature of the file. As with this example, where there is a large amount of mean drift. If multiple segments have been removed in a row, ***flowCut*** will extend this removal section until the bordering segment has less than or equal to 1 standard deviation away from the mean of the means of the segments still not removed. In other words, segments to the far right were first selected as significantly different and marked for removal. However, ***flowCut*** then extended its range to the left until the brown mean line showed a more flattened behaviour throughout the file.

To convince you and to illustrate this in a different way, we can compare the results of setting *GateLineForce* to 44 (shown in Figure 7) with our previous result from Figure 6:

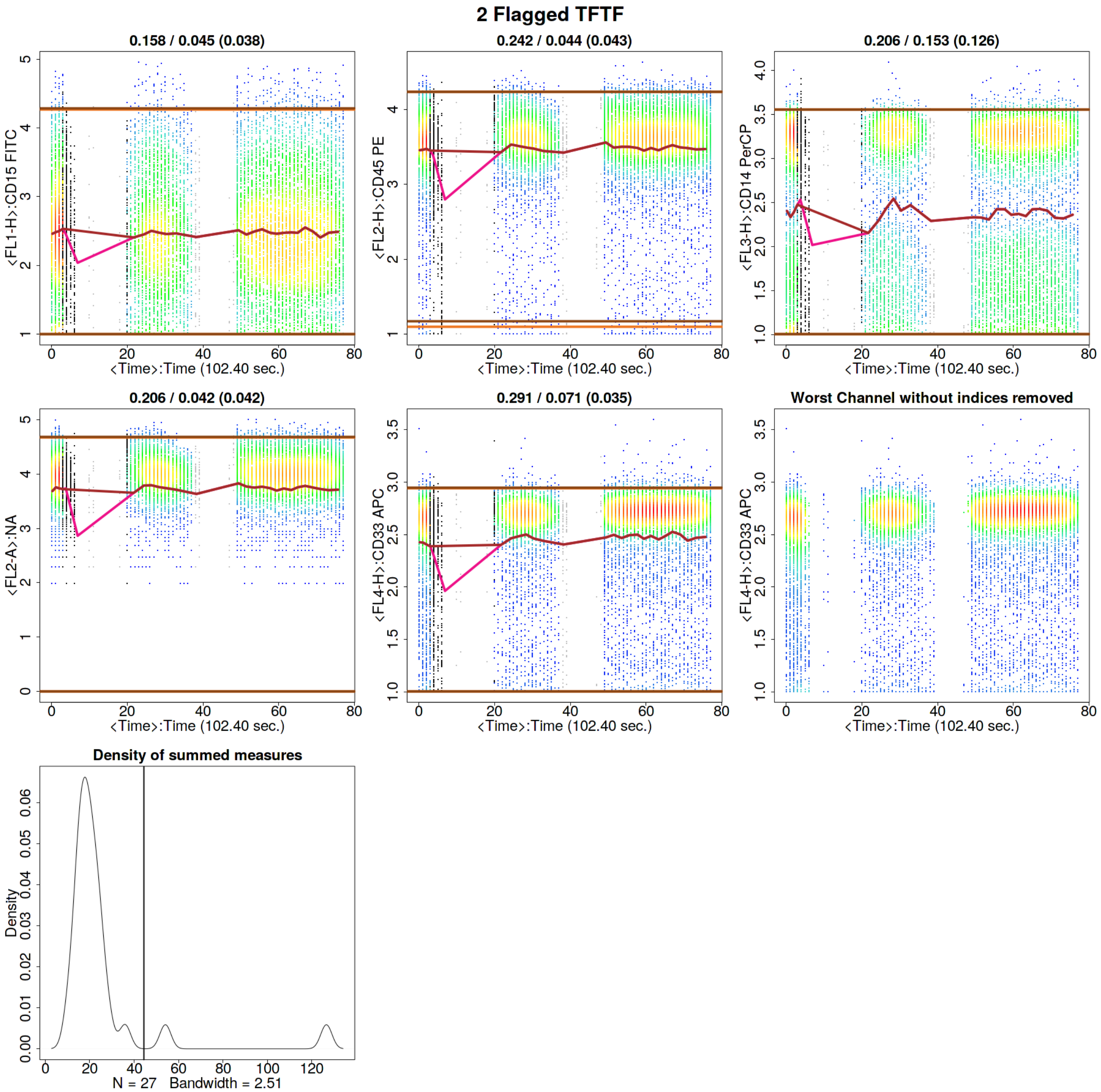


Figure 3: Running ***flowCut*** with *MaxValleyHgt* changed to 0.01.

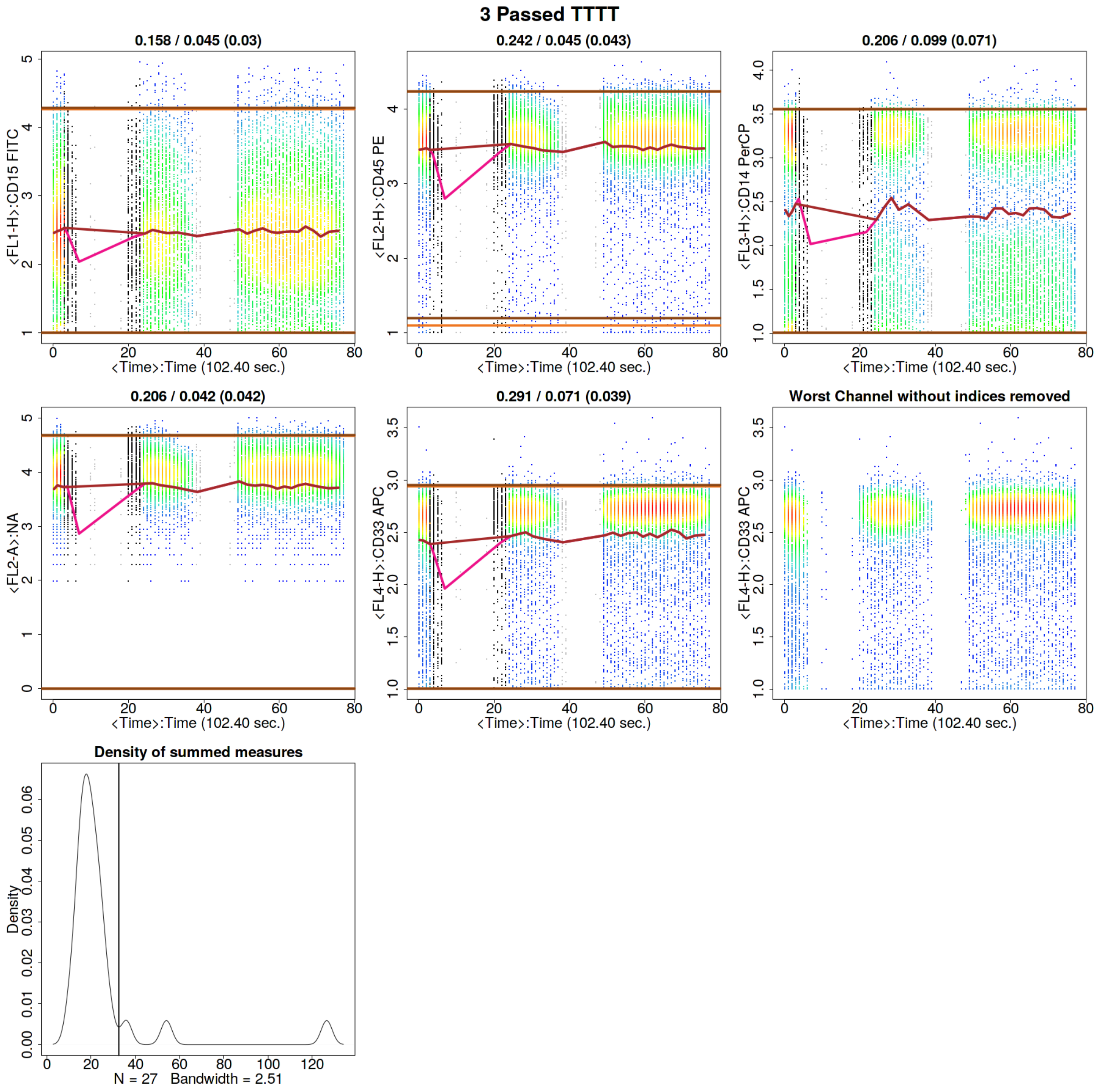


Figure 4: Running ***flowCut*** with default *MaxValleyHgt*.

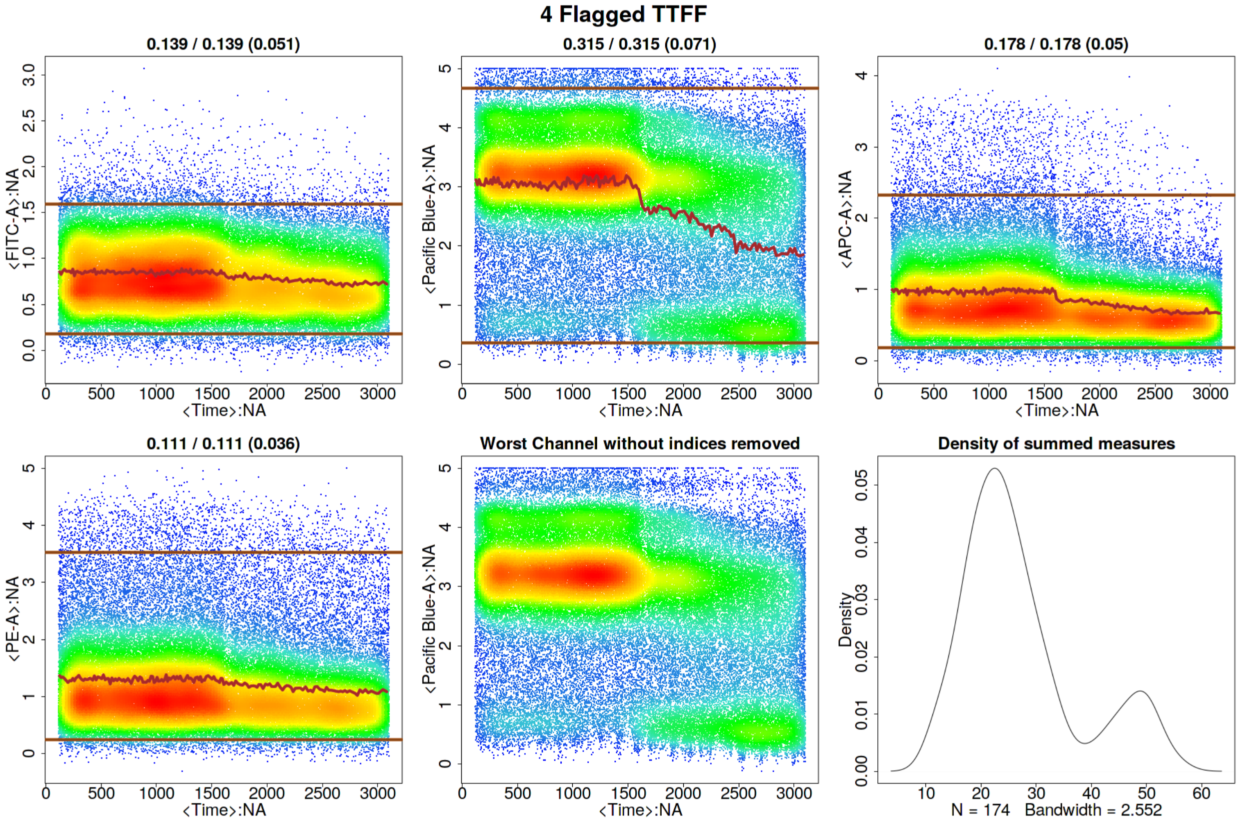


Figure 5: Running ***flowCut*** with *MaxPercCut* changed to 0.15.

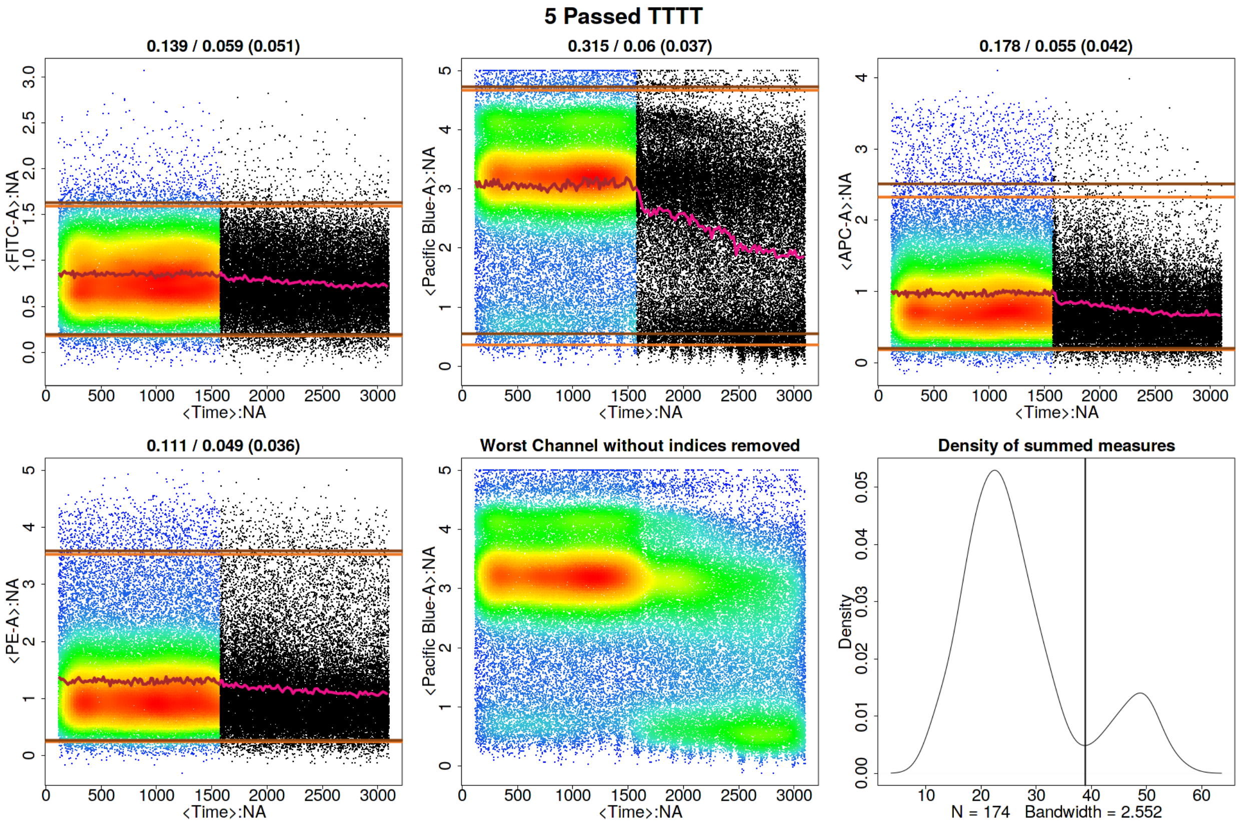


Figure 6: Running ***flowCut*** with default *MaxPercCut*.

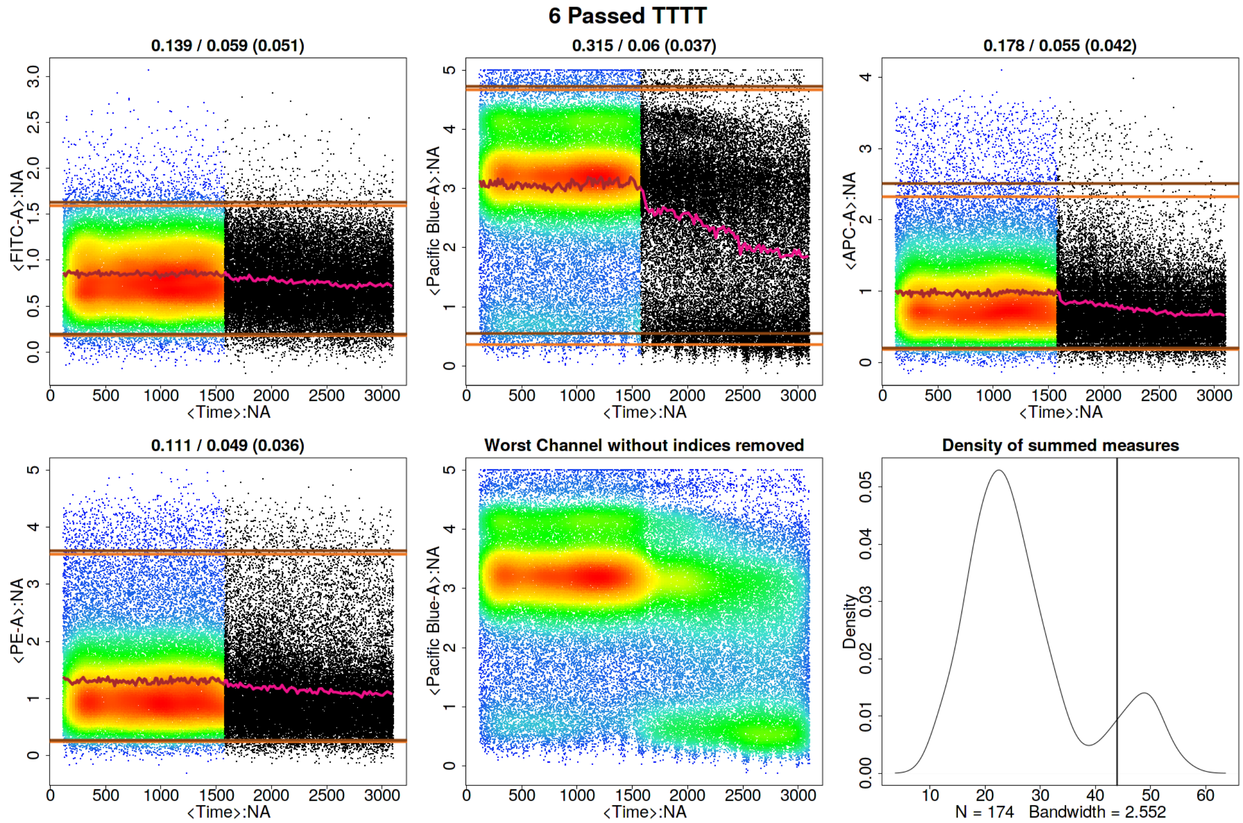


Figure 7: Running ***flowCut*** with *GateLineForce* set to 44.

```
res_flowCut <- flowCut(flowCutData[[3]], FileID=6, Plot ="All", GateLineForce=44)
```

The intention of *GateLineForce* is to give the user the option to override the automatedly calculated cut off point between segments to keep and segments to remove. This is intended as a last option for the user. In this case we have chosen *GateLineForce* to be 44 to show a particular behaviour of flowCut that allows additional removal of segments. Therefore, even though the gating threshold has changed we still have the same amount of events removed.

```
res_flowCut$data["% of events removed",]
```

```
## % of events removed
##              "42.27"
```

This happened because in both cases there was a large section in a row removed and ***flowCut*** expanded the section until all that was left was not too variant.

#### Parameters *UseOnlyWorstChannels* and *AllowFlaggedRerun*

Sometimes abnormal behaviours are only present in a few channels. The outlier events in these channels can be washed out when taking into consideration all measures in all channels. As a result, outliers in these channels are not properly removed. In these cases, we can use the parameter *UseOnlyWorstChannels* to only focus on these specific channels.

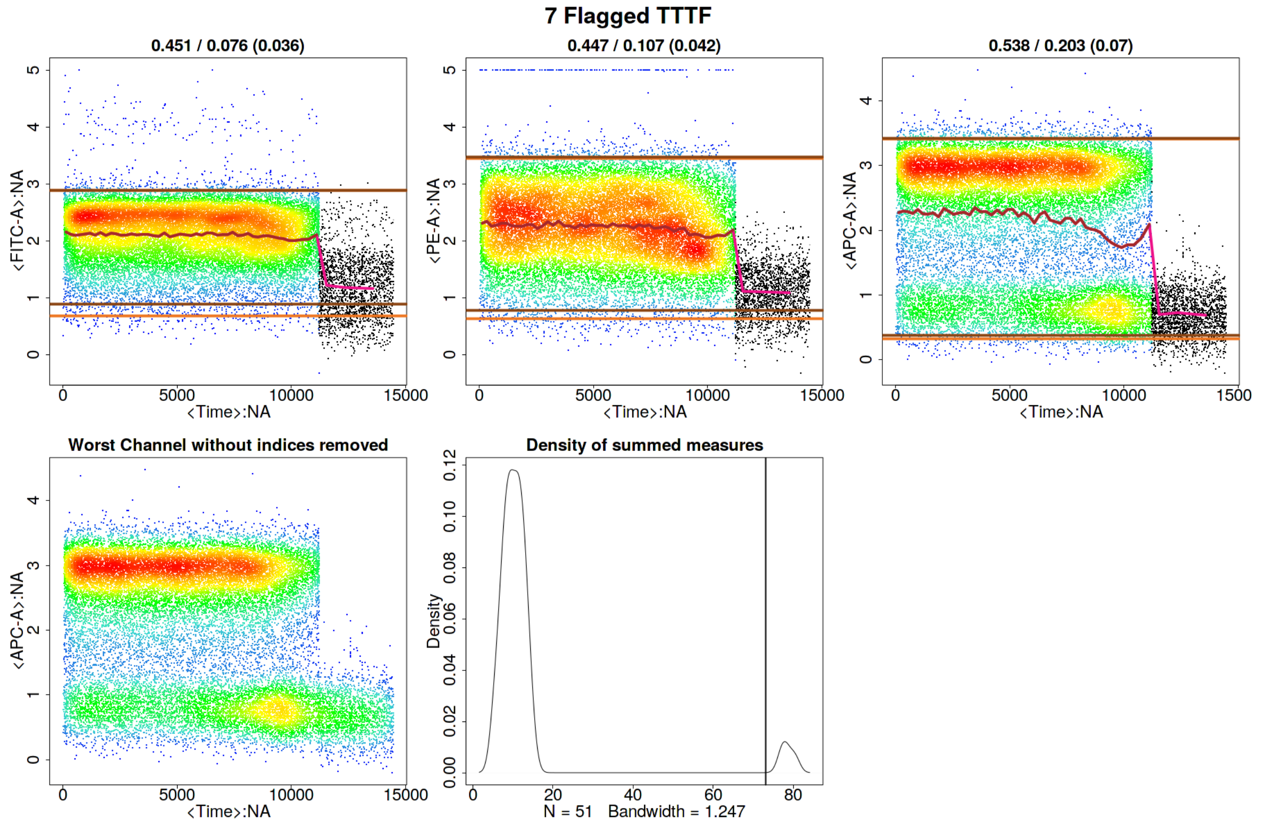


Figure 8: Running ***flowCut*** with the default values.

In other cases, some files may contain two or more groups of outlier events, where one group of outliers is significantly different than the others. The first round of ***flowCut*** will remove the most significantly different outlier groups while other outlier groups are masked by the worst ones. This issue can be addressed by using the parameter *AllowFlaggedRerun*, which will rerun the algorithm a second time if it has been flagged the first time.

An example of when both the above parameters are required is shown in Figure 8, but first showing results of ***flowCut*** with default values.

```
res_flowCut <- flowCut(flowCutData[[4]], FileID=7, Plot ="All")
```

The section with time greater than approximately 11000 is drastically different than the rest of the file, however, the section with time between 7500 and 11000 is still different enough and should be removed. The most deviant population of events have been removed. But, the file is still flagged because the third channel shows a mean drift that's more than the defaulted value for *MaxOfMeans*. It would be a good idea to run ***flowCut*** again on this file to remove the mean drift events. One thing to be noted here is that the mean drift occurs strongly in only one channel. To properly catch those events and to counter the averaging effect, we need to set the *UseOnlyWorstChannels* parameter to 'TRUE'. The result of the second round of cleaning and using only the worst channels is shown in Figure 9. Please note the asterisks in the figure, which show which channels were selected as the worst channels.

```
res_flowCut <- flowCut(flowCutData[[4]], FileID=8, AllowFlaggedRerun = TRUE,
                       UseOnlyWorstChannels = TRUE, Plot="All")
```

In this example two channels were selected as the worst channels. The parameters *AmountMeanRangeKeep*

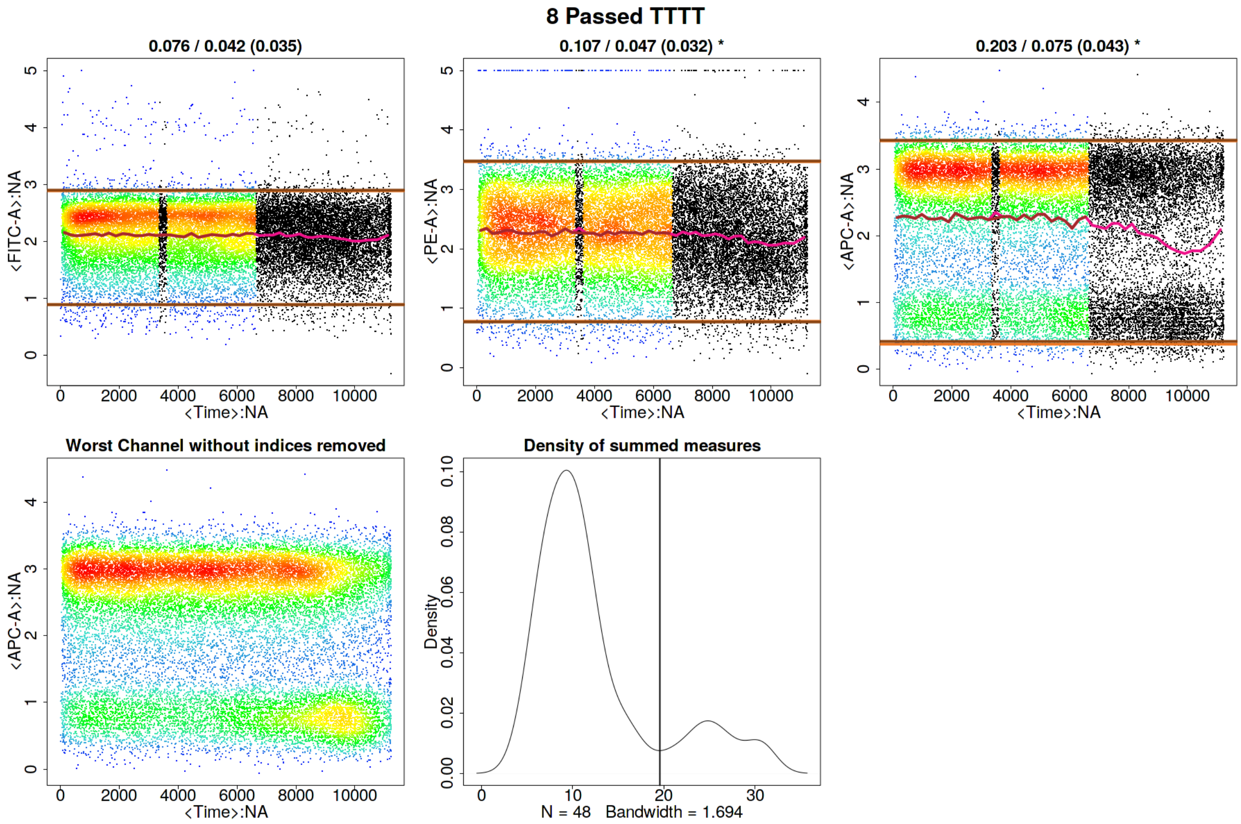


Figure 9: Running ***flowCut*** a second time using only the worst channels.

and *AmountMeanSDKeep*, defaulted at 1 and 2 respectively, determine how many channels of each type of worst channel detection are chosen as a worst channel. *AmountMeanRangeKeep* looks for the channels that have the highest range in their segments' means. *AmountMeanSDKeep* looks for the channels that have the highest variance (SD) in their segments' means. With the defaults, ***flowCut*** will return either 3 worst channels if there is no overlap or 2 worst channels if there is overlap, as there is in the above example. Please note that when using *UseOnlyWorstChannels*, it will calculate the worst channels independently for each file.

#### ***Segment*** Size

The size of *Segment* can significantly impact the result of ***flowCut***. The algorithm groups 500 events together in a segment and compares the measures of the segment against all other segments. The number of 500 has proven successful on almost all files. We advise not to lower or increase this number unless there are specific reasons to do so. Without enough events in each segment, statistics start becoming less meaningful. 500 is small enough to catch spikes from clogs but large enough for good statistics. And spikes that take up 1000 events in a 1 000 000 event file do not have analogous spikes of 10 events in a 10 000 event file. The spikes will still be 1000 events in a 10 000 event file. With that being said, there are cases that it makes sense to change the segment size.

Depending on the size of the file, there is an optimal amount of events to be grouped together for the best performance of the outlier detection. Users should ensure there are at least 100 segments in total (total event count divided by *Segment*) for analysis for small files. For large files, users could increase the *Segment* parameter if large fluctuation in the mean is causing an inaccurate detection of outliers (i.e. if the brown line is too variable, then increasing the *Segment* parameter may be beneficial). For example, we analyzed a 65MB file (hence data not attached) with the *Segment* parameter equal to 500. Figure 10 shows large fluctuations in the means and random segment deletions without catching the mean drift. However, if we increase *Segment* to 5000, the algorithm can correctly detect and remove the part of the events that are deviant. The result is shown in Figure 11.

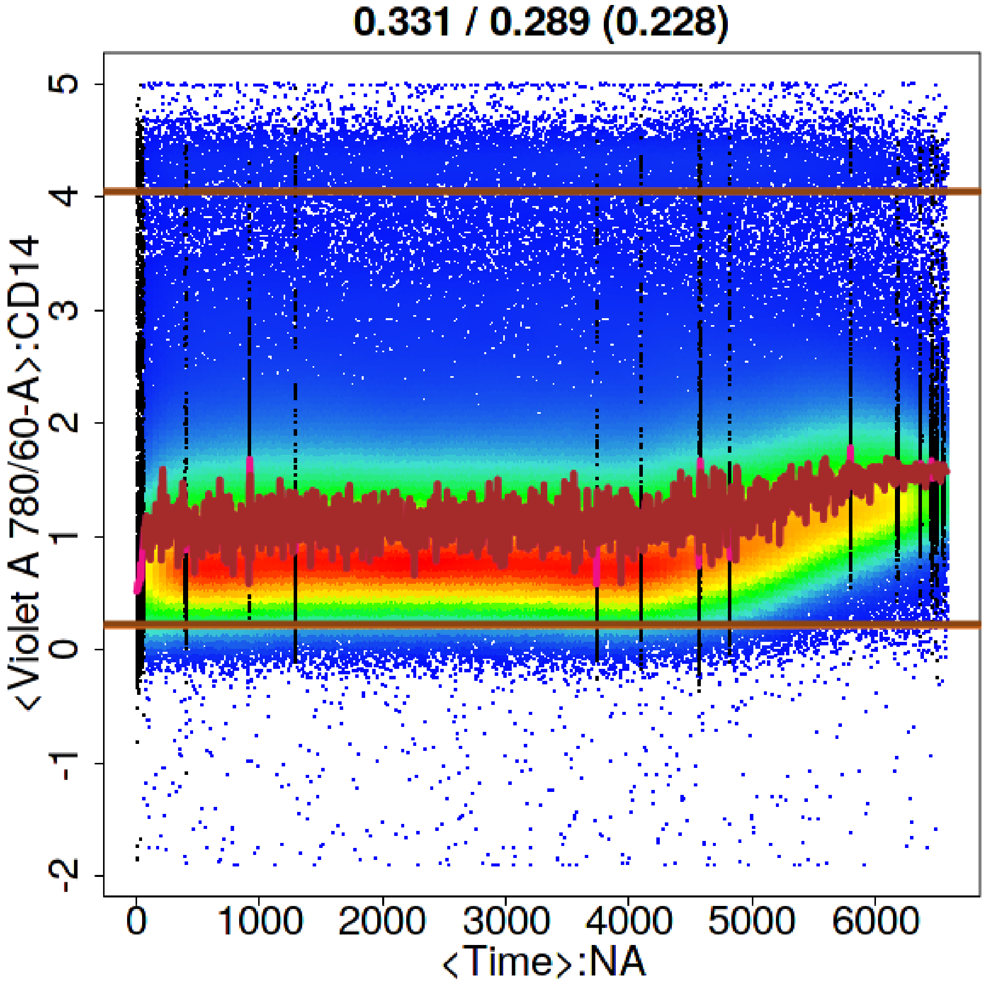


Figure 10: Running ***flowCut*** with *Segment* set to 500.

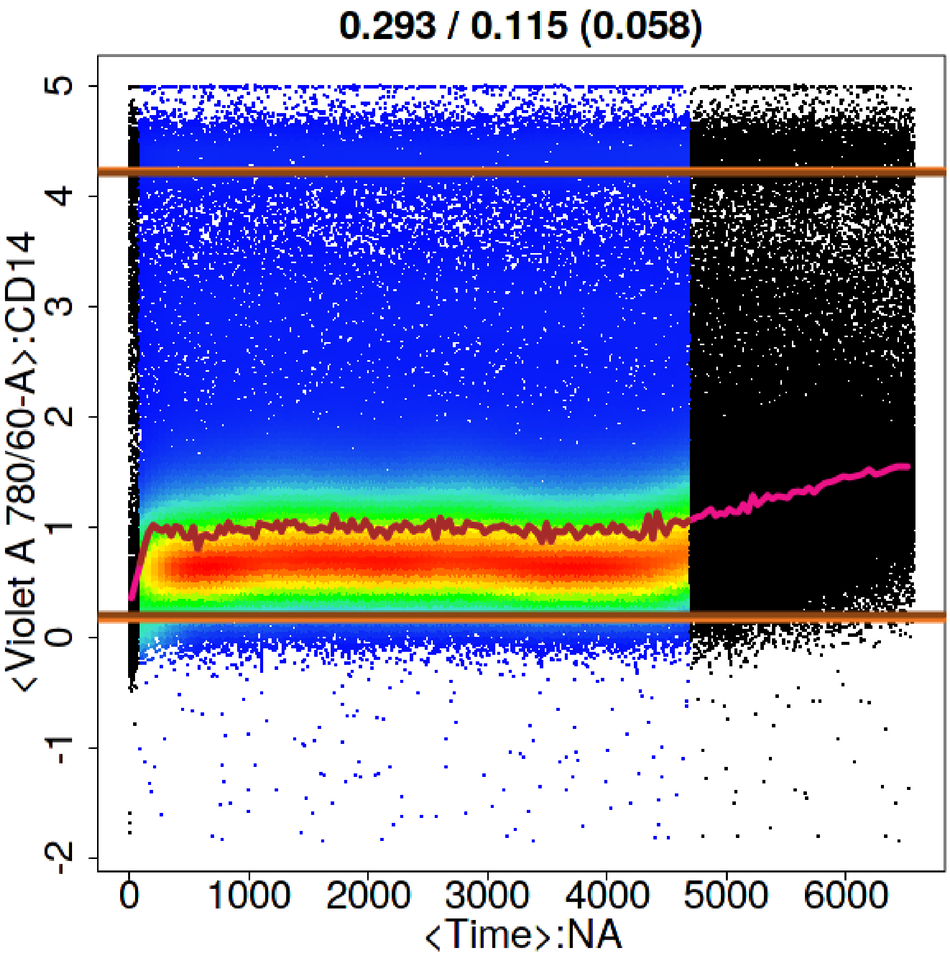


Figure 11: Running ***flowCut*** with *Segment* set to 5000.
