## Supplementary material for "flowCut — An R package for precise and accurate automated removal of outlier events and flagging of files based on time versus fluorescence analysis": paper_code: F1_byproblemtype_calculation.R

```
library(ggplot2)
library(tidyverse)

F1.score.table <- read.csv(file=paste0("~/results/55/F1_scores_results_byProblemType.csv"))

F1.score.table[which(F1.score.table[,"Type"]=="spke"),"Type"] <- "spike"

#bar plot
F1.score.table$Type <- as.factor(F1.score.table$Type)
bar.plot.data <- aggregate(F1.score.table[,c(4,7,10)], list(F1.score.table$Type),mean)
bar.plot.data <- bar.plot.data %>% gather(type,score,F1_fCut:F1_fClean) 
print(bar.plot.data) #show F1 scores by problem type
bar.plot.data$type <- as.factor(bar.plot.data$type)
bar.plot.data$Group.1 <- factor(bar.plot.data$Group.1,levels = c("shift","spike","low_dens",
                                                           "discontinuity","clean"))
ggplot(bar.plot.data,aes(fill=type,y=score,x=Group.1)) +
geom_bar(position="dodge", stat="identity") + theme_bw()


  pie.chart.data <- F1.score.table %>% group_by(Type) %>% summarize(freq=n())

  pie.chart.data <- c(as.character(unique(F1.score.table$Type)))  
  pie.chart.data <- pie.chart.data[-which(pie.chart.data=="clean")]
  
  counts.pie <- NULL
  for(n in 1:length(pie.chart.data)){
    counts.pie <- c(counts.pie, length(unique(F1.score.table$Name[which(F1.score.table$Type == pie.chart.data[n])])))
  }  
  
  pie.chart.data <- data.frame(pie.chart.data,counts.pie)
  colnames(pie.chart.data) <- c("problem_type","file_counts")
  pie.chart.data$problem_type <- as.factor(pie.chart.data$problem_type)
  
  #bar plot #2
  bp2.data <- aggregate(F1.score.table[,c(13)], list(F1.score.table$Type),sum)
  bp2.data[,2] <- round((bp2.data[,2] / c(52,3,12,28,18)) *100,2)
  
  bp2.data <- bp2.data[-1,]
  colnames(bp2.data) <- c("problem_type","Percentage")
                                                         
  bp3.data <- left_join(bp2.data,pie.chart.data)
  bp3.data[,2] <-  round(bp3.data[,2] * bp3.data[,3] /52,2)
  
  #bp3 shows percentage of each problem type in a file
  bp3 <- ggplot(bp3.data,aes(y=Percentage,x=reorder(problem_type,-Percentage))) +
    geom_bar(position="dodge", stat="identity",width=0.7,fill="black", color="black") +
    geom_text(aes(label= paste0(Percentage," %")), vjust=-0.5 )+
    scale_y_continuous(expand = c(0,0),
                       limits = c(0,30))+
    theme_bw() + xlab("") +
    ylab("") + theme(axis.text=element_text(size=13),
                     axis.text.y = element_blank(),
                     panel.grid.major = element_blank(), 
                     panel.border = element_blank(),
                     panel.grid.minor = element_blank(),
                     axis.ticks = element_blank())
  
  ggsave(bp3,filename = paste0("~/results/bar2.pdf"))
  

 
```
