## Supplementary material for "flowCut — An R package for precise and accurate automated removal of outlier events and flagging of files based on time versus fluorescence analysis": paper_code: flowCut_Cleaning_Comparison.R

```
#############################################################################
# This code is to accompany the flowCut Paper published with Bioinformatics #
# Title:                                                                    #
## flowCut — An R package for precise and accurate                          #
## automated removal of outlier events and flagging                         #
## of files based on time versus fluorescence analysis                      #
# Written by Justin Meskas and Sherrie Wang                                 #
# Last Updated September 2018                                               #
#############################################################################

library("flowCore") # 1.51.7
library("flowDensity") # 1.19.5
library("flowCut") # 0.99.26
library("flowClean") # 1.24.0
library("flowAI") # 1.18.1
library("doMC") # 1.3.6
library("foreach") # 1.4.7
library("Cairo") # 1.5-10
library("FlowRepositoryR") # 1.14.1
require("R.utils") # 2.9.0
library("stringr") # 1.4.0

#setwd("/code/Projects/flowCut_Paper/codes_plus/")
source("code/flowCut_Cleaning_Comparison_Functions.R")

start1 <- Sys.time()
set.seed(5678) # flowAI

no_cores <- detectCores() - 1
registerDoMC(no_cores)

skip_flowClean <- F
skip_flowAI <- F
skip_Plotting <- T

flowCutFailed <- NULL
flowCleanFailed <- NULL
flowAIFailed <- NULL

flowCutTimeout <- NULL
flowCleanTimeout <- NULL
flowAITimeout <- NULL

# download files from FlowRepository
suppressWarnings ( dir.create ( paste0(getwd(), "/data/FlowRepFiles/"), recursive = T ))
if ( length(dir(paste0(getwd(), "/FlowRepFiles/"), pattern = ".fcs")) != 55 ) {
    myDatasets <- flowRep.get("FR-FCM-ZYPD")
    sapply(1:length(fcs.files(myDatasets)), function(x) { download(fcs.files(myDatasets)[[x]], dirpath=paste0(getwd(), "/data/FlowRepFiles/"), show.progress=F)})
    sapply(1:length(myDatasets@attachments), function(x) { download((myDatasets@attachments)[[x]], dirpath=paste0(getwd(), "/data/FlowRepFiles/"), show.progress=F)})
}

suppressWarnings ( dir.create ( paste0(getwd(), "/results/55/"), recursive = T ))

# Randomly selected flowCut Repository True Positive files
fileNames <- dir(paste0(getwd(), "/data/FlowRepFiles"), pattern = ".fcs", recursive = T, full.names = F)
allFiles <- dir(paste0(getwd(), "/data/FlowRepFiles"), pattern = ".fcs", recursive = T, full.names = T)


# Subcategorize 55 files, Category one is for files with clear visual boundaries for removal
# Category two is files that are obviously problems, but the boundary for removal is fuzzy.
# Category three is files that are either obviously wrong but don't know which part to remove that can 'fix' the files,
# or the removal regions are not obvious to the manual gater, but are identified by all three algorithms, however, the 
# regions are not necesarilly overlapping, creating confusion as to which are the true positive regions. 
# Category three files are removed from F1 score calculation as there is no gold standard.
# Category three files are there to examine how each algorithm deal with files like this.
category <- 2

# read in manual gating data
gating_thresholds <- unname(as.matrix(read.csv(paste0(getwd(), "/data/FlowRepFiles/Manual_Gating_52.csv"))))


if(category == 1){
  fileNames <- dir(paste0(getwd(), "/data/FlowRepFiles/Category_One"), pattern = ".fcs", recursive = T, full.names = F)
  allFiles <- dir(paste0(getwd(), "/data/FlowRepFiles/Category_One"), pattern = ".fcs", recursive = T, full.names = T)
  cat.ind <- sapply(fileNames,function(fn)which( gating_thresholds[,1] == fn))
  gating_thresholds <- gating_thresholds[cat.ind,]
}
if(category == 2){
  fileNames <- dir(paste0(getwd(), "/data/FlowRepFiles/Category_Two"), pattern = ".fcs", recursive = T, full.names = F)
  allFiles <- dir(paste0(getwd(), "/data/FlowRepFiles/Category_Two"), pattern = ".fcs", recursive = T, full.names = T)
  cat.ind <- sapply(fileNames,function(fn)which( gating_thresholds[,1] == fn))
  gating_thresholds <- gating_thresholds[cat.ind,]
}
if(category == 3){#for cases when there are no gold standard.
  fileNames <- dir(paste0(getwd(), "/data/FlowRepFiles/Category_Three"), pattern = ".fcs", recursive = T, full.names = F)
  allFiles <- dir(paste0(getwd(), "/data/FlowRepFiles/Category_Three"), pattern = ".fcs", recursive = T, full.names = T)
  cat.ind <- sapply(fileNames,function(fn)which( gating_thresholds[,1] == fn))
  gating_thresholds <- gating_thresholds[cat.ind,]

}

######## Begin foreach ########
results <- foreach (q1 = c(1:length(fileNames)), .combine = rbind, .maxcombine = length(fileNames), .multicombine = T) %dopar% {
# results <- foreach (q1 = 3:5, .combine = rbind, .maxcombine = 3, .multicombine = T) %dopar% {

        precision_fAI <- 0
        precision_fcut <- 0
        precision_fclean <- 0
        recall_fAI <- 0
        recall_fcut <- 0
        recall_fclean <- 0
        F1_fcut <- 0
        F1_fAI <- 0
        F1_fclean <- 0

        x1 <- which(fileNames == gating_thresholds[q1,1])

        manual_gates <- as.numeric(gating_thresholds[q1,2:ncol(gating_thresholds)])
        manual_gates <- manual_gates[!is.na(manual_gates)]

        fileId <- fileNames[x1]
        print(paste0("Starting ", fileNames[x1]))
        fz <- try (read.FCS(allFiles[x1]))
        if(class(fz)=="try-error"){
            index.Corrupted <- "Corrupted"
            Desc <- Name <- NumEvents <- lessThan20000 <- ind.clean <- numClean <- res.flowCut.data <- NULL
            if(!file.exists("filecorrupt.txt")){
                write.table(t(c("file Corrupt :", fileId, fileNames[x1])),col.names = FALSE, file = "results/55/filecorrupt.txt")
            }
            write.table(t(c("file Corrupt :", fileId, fileNames[x1])),col.names = FALSE, file = "results/55/filecorrupt.txt", append = TRUE)

        } else {
            index.Corrupted <- "NotCorrupt"
            Desc <- fz@parameters@data$desc
            Name <- fz@parameters@data$name
            NumEvents <- nrow(fz)
            f_org <- fz
            if ( nrow(fz) <= 2000){
                lessThan20000 <- "less2k"
            } else {
                lessThan20000 <- "more2k"
            }

            scat.chans <- c(grep (colnames(fz),pattern = "FSC*"),grep (colnames(fz),pattern = "SSC*"))
            if(length(scat.chans)==0){
              f.comp <- fz
            }else{
              f.marg <- removeMargNegs(fz, chans=scat.chans, neg=-1, verbose= F)
              f.comp <- f.marg$f; ind.marg.neg <- f.marg$ind.marg.neg; remove(f.marg)
            }

            ######## Compensation ########
            des <- fz@description
            matrix_ind <- grep(pattern = "spill", x=names(des),ignore.case = T)
            if(length(matrix_ind)==0){
                message("no spill over matrix is found, skipping compensation...")
            } else{

                if ( det(des[matrix_ind][[1]])==1 ) {
                    message("Check the spillover matrix, it's probably an identity matrix!")
                } else {
                    f.comp <- compensate(f.comp, des[matrix_ind][[1]])
                }

            }
            ######## Transformation ########
            time.loc <- which(tolower(f.comp@parameters@data$name) == "time"); names(time.loc) <- NULL
            channels.ind <- setdiff(1:length(colnames(f.comp)), c(scat.chans, time.loc)) #locate channels other than scatter channels

            lgl <- try (estimateLogicle(f.comp, channels= colnames(f.comp)[channels.ind], m=5))
            if(class(lgl)=="try-error"){

                if(!file.exists("filecorrupt.txt")){
                    write.table(t(c("file not standard :", fileId, fileNames[x1])),col.names = FALSE, file = "results/55/filecorrupt.txt")
                }
                write.table(t(c("file not standard :", fileId, fileNames[x1])), col.names = FALSE, file = "results/55/filecorrupt.txt", append = TRUE)
                {next}
            }
            fs.trans <- transform(f.comp, lgl)

            z <- ceiling(sqrt(length(channels.ind)+2))
            # suppressWarnings ( dir.create ( paste0(getwd(), "/results/55/flowCutRep_Plots/"), recursive = T ))
            # CairoPNG ( file = paste0(getwd(), "/results/55/flowCutRep_Plots/", fileNames[x1],".png"), height = z*600, width = z*600)
            # 
            # par(mfrow=c(z,z),mar=c(7,7,4,2),mgp=c(4,1.5,0))
            # 
            # for (x in 1:length(channels.ind)){
            # plotDens(fs.trans, c(time.loc,channels.ind[x]), cex.main=3, cex.lab=3, cex.axis=3,
            #          main=paste0(fileNames[x1],"_",x))
            # }
            # dev.off()

            ######## Manual Gating ########
            toremoveRegion <- NULL
            manual_gates <- split(manual_gates, ceiling(seq_along(manual_gates)/2))
            manual_gates <- unname(manual_gates)
            for (gs in 1:length(manual_gates)){
              a  <- which(fs.trans[,time.loc]>manual_gates[[gs]][1])
              a1 <- which(fs.trans[,time.loc]<manual_gates[[gs]][2])
              toremoveRegion <- c(toremoveRegion, intersect(a, a1))
            }

            ######## random cleaning ########
            removeRegion_Rand <- sample(x=1:nrow(fs.trans), length(toremoveRegion))
            precision_rand <- length(intersect(removeRegion_Rand,toremoveRegion))/length(toremoveRegion)
            if(is.nan(precision_rand)){ precision_rand <- 0 }
            recall_rand <- length(intersect(removeRegion_Rand,toremoveRegion))/length(removeRegion_Rand)
            if(is.nan(recall_rand)){ recall_rand <- 0 }
            F1_rand <- 2*(precision_rand*recall_rand)/(precision_rand+recall_rand)
            if(is.nan(F1_rand)){ F1_rand <- 0 }

            ######## flowCut ########
            start.flowCut <- Sys.time()
            print("Starting flowCut")
            segment <- 500

            res.flowCut <- try( flowCut(f=fs.trans, Segment=segment, FileID = fileId, Directory = paste0(getwd(),"/results/55/flowCutRep_Plots/"),
                                        Plot ="None",AllowFlaggedRerun =  TRUE, Verbose=TRUE) )

            if (class(res.flowCut)=="try-error"){
                resTable <- NULL
                f.tvsf <- fs.trans
                ind.removed <- NULL
                flowCutFailed <- c(flowCutFailed, x1)
                if(!file.exists("results/55/flowCutError.txt")){
                  write.table(t(c("flowCut Error File :", fileId)),col.names = FALSE, file = "results/55/flowCutError.txt")
                }
                write.table(t(c("flowCut Error File :", fileId)),col.names = FALSE, file = "results/55/flowCutError.txt", append = TRUE)

            } else{
                resTable <- res.flowCut$data
                f.tvsf <- res.flowCut$frame
                ind.removed <- res.flowCut$ind
                if(length(ind.removed)==0){
                    ind.removed <- 0
                }
                precision_fcut <- length(intersect(ind.removed,toremoveRegion))/length(toremoveRegion)
                if(is.nan(precision_fcut)){ precision_fcut <- 0 }
                recall_fcut <- length(intersect(ind.removed,toremoveRegion))/length(ind.removed)
                if(is.nan(recall_fcut)){ recall_fcut <- 0 }
                F1_fcut <- 2*(precision_fcut*recall_fcut)/(precision_fcut+recall_fcut)
                if(is.nan(F1_fcut)){ F1_fcut <- 0 }

                Time.flowCut <- TimeOutput(start.flowCut)
            }

            ######## flowAI ########
            if (skip_flowAI == F) {
                print("Starting flowAI")
                start.flowAI <- Sys.time()
                flowAIError <- ""

                    res.flowAI <- withTimeout({
                        try.flowAI <- try(fAI.cleaned_All <- flow_auto_qc(fs.trans, html_report = FALSE, mini_report = FALSE,
                                                                          remove_from="FS_FM", second_fractionFR = 0.1, fcs_QC = FALSE, output = 2 ))

                        if(class(try.flowAI) =="try-error"){
                            fAI.cleaned2_All <- fs.trans
                            to.be.removed.AI_All <- NULL
                            flowAIError <- "Error"
                            flowAIFailed <- c(flowAIFailed, paste0(fileId, "_", x1))
                            cat(paste0("flowAI with the current values does not work.", "\n"))
                            if(!file.exists("results/55/flowAIError.txt")){
                              write.table(t(c("flowAI Error File :", fileId)),col.names = FALSE, file = "results/55/flowAIError.txt")
                            }
                            write.table(t(c("flowAI Error File :", fileId)),col.names = FALSE, file = "results/55/flowAIError.txt", append = TRUE)
                        } else {
                            rem.ind_All <- which(fAI.cleaned_All@parameters@data$name == "remove_from_FS_FM")
                            to.be.removed.AI_All <- which(fAI.cleaned_All@exprs[,rem.ind_All] >= 9999)
                            fAI.cleaned2_All <- fAI.cleaned_All
                            if( length(to.be.removed.AI_All) >= 1 ){
                                fAI.cleaned2_All@exprs <- fAI.cleaned_All@exprs[-to.be.removed.AI_All,]
                                precision_fAI <- length(intersect(to.be.removed.AI_All,toremoveRegion))/length(toremoveRegion)
                                if(is.nan(precision_fAI)){ precision_fAI <- 0 }
                                recall_fAI <- length(intersect(to.be.removed.AI_All,toremoveRegion))/length(to.be.removed.AI_All)
                                if(is.nan(recall_fAI)){ recall_fAI <- 0 }
                                F1_fAI <- 2*(precision_fAI*recall_fAI)/(precision_fAI+recall_fAI)
                                if(is.nan(F1_fAI)){ F1_fAI <- 0 }
                            }
                        }
                    }, timeout=3600);

                Time.flowAI <- TimeOutput(start.flowAI)
                fAI.cleaned3_All <- fAI.cleaned2_All
                fAI.cleaned3_All@exprs[,time.loc] <- 1:nrow(fAI.cleaned3_All)
            }

            ######## flowClean ########
            if (skip_flowClean == F) {
                start.flowClean <- Sys.time()
                print("Starting flowClean")
                FSC.loc  <- grep("fsc", tolower(fs.trans@parameters@data$name));  names( FSC.loc) <- NULL
                SSC.loc  <- grep("ssc", tolower(fs.trans@parameters@data$name)); names( SSC.loc) <- NULL
                Time.loc <- which(tolower(fs.trans@parameters@data$name) == "time"); names(Time.loc) <- NULL
                CleanChan.loc <- (1:ncol(fs.trans))[-c(FSC.loc, SSC.loc, Time.loc)]
                flowCleanError <- ""
                    res.fclean <- withTimeout({
                        try.flowClean <- try(f.clean <- clean(fs.trans, vectMarkers=CleanChan.loc, ext="fcs", filePrefixWithDir=paste("temp/Clean",sep="/")))
                        if(class(try.flowClean) =="try-error"){
                            f.clean <- fs.trans
                            to.be.removed.fC <- NULL
                            flowCleanError <- "Error"
                            flowCleanFailed <- c(flowCleanFailed, paste0(fileId, "_", x1))
                            cat("flowClean with the current values does not work. \n")
                            if(!file.exists("results/55/flowCleanError.txt")){
                              write.table(t(c("flowClean Error File :", fileId)),col.names = FALSE, file = "results/55/flowCleanError.txt")
                            }
                            write.table(t(c("flowClean Error File :", fileId)),col.names = FALSE, file = "results/55/flowCleanError.txt", append = TRUE)
                        } else {
                            rg <- rectangleGate(filterId="gvb", list("GoodVsBad"=c(0, 9999)))
                            idx <- flowCore::filter(f.clean, rg)
                            if(length(which(idx@subSet==F))!=0){
                                to.be.removed.fC <- which(idx@subSet==F)
                                exprs(f.clean) <- exprs(f.clean)[-which(idx@subSet==F),]
                                precision_fclean <- length(intersect(to.be.removed.fC,toremoveRegion))/length(toremoveRegion)
                                if(is.nan(precision_fclean)){ precision_fclean <- 0 }
                                recall_fclean <- length(intersect(to.be.removed.fC,toremoveRegion))/length(to.be.removed.fC)
                                if(is.nan(recall_fclean)){ recall_fclean <- 0 }
                                F1_fclean <- 2*(precision_fclean*recall_fclean)/(precision_fclean+recall_fclean)
                                if(is.nan(F1_fclean)){ F1_fclean <- 0 }
                            } else {
                                f.clean <- fs.trans
                                to.be.removed.fC <- NULL
                            }
                        }
                    }, timeout=3600*6);
                Time.flowClean <- TimeOutput(start.flowClean)
                f.clean3 <- f.clean
                f.clean3@exprs[,time.loc] <- 1:nrow(f.clean3)
            }

            ######## plot figures ########
            if (skip_Plotting == F) {
                x <- res.flowCut$worstChan
                suppressWarnings ( dir.create ( paste0(getwd(), "/results/55/Comparison/"), recursive = T ))

                HowManyMore <- max(0, length(channels.ind)-1)
                if (is.null(x)){
                    x <- sample(channels.ind,1)
                }
                channels.ind.refined <- c(x, sample(channels.ind[-which(channels.ind == x)], HowManyMore))
                ChannelsToPlot <- fs.trans@parameters@data$desc[channels.ind.refined]
                widthUnits <- length(ChannelsToPlot)
                CairoPNG ( file = paste0(getwd(), "/results/55/Comparison/", resTable["FileID",], "_", segment, ".png"),
                           height = 5*400, width = widthUnits*400)

                par(mfrow=c(5,widthUnits),mar=c(5,5,4,2))

                for ( e1 in 1:widthUnits){
                    plotDens(fs.trans, c(time.loc,channels.ind.refined[e1]), cex.main=2, cex.lab=2, cex.axis=2, main=paste0("No events removed - ", ChannelsToPlot[e1]))
                    # points(exprs(fs.trans)[toremoveRegion ,c(time.loc,channels.ind.refined[e1])],pch=".",col="black")
                }

                for ( e1 in 1:widthUnits){
                    plotDens(fs.trans, c(time.loc,channels.ind.refined[e1]), cex.main=2, cex.lab=2, cex.axis=2, main=paste0("Manual Results - ", ChannelsToPlot[e1]))
                    points(exprs(fs.trans)[toremoveRegion ,c(time.loc,channels.ind.refined[e1])],pch=".",col="black")
                }

                for ( e1 in 1:widthUnits){
                    plotDens(fs.trans, c(time.loc,channels.ind.refined[e1]), cex.main=2, cex.lab=2, cex.axis=2, main=paste0("flowCut - ", ChannelsToPlot[e1]))
                    points(exprs(fs.trans)[res.flowCut$ind ,c(time.loc,channels.ind.refined[e1])],pch=".",col="black")
                }

                for ( e1 in 1:widthUnits){
                    plotDens(fs.trans, c(time.loc,channels.ind.refined[e1]), cex.main=2, cex.lab=2, cex.axis=2, main=paste0("flowAI - ", ChannelsToPlot[e1]))
                    points(exprs(fs.trans)[to.be.removed.AI_All,c(time.loc,channels.ind.refined[e1])],pch=".",col="black")
                }

                for ( e1 in 1:widthUnits){
                    plotDens(fs.trans, c(time.loc,channels.ind.refined[e1]), cex.main=2, cex.lab=2, cex.axis=2, main=paste0("flowClean - ", ChannelsToPlot[e1]))
                    points(exprs(fs.trans)[to.be.removed.fC ,c(time.loc,channels.ind.refined[e1])],pch=".",col="black")
                }
                dev.off()
            }

            ######## calculate events removed ########
            Manu_rem <- round(length(toremoveRegion)/length(fs.trans@exprs[,1]) *100, digits = 3)
            Rand_rem <- round(length(removeRegion_Rand)/length(fs.trans@exprs[,1]) *100, digits = 3)
            fCut_rem <- round((length(fs.trans@exprs[,1])-length(f.tvsf@exprs[,1]))/length(fs.trans@exprs[,1]) *100, digits = 3)
            if (skip_flowAI == F){
                fAI_rem <- round((length(fs.trans@exprs[,1])-length(fAI.cleaned2_All@exprs[,1]))/length(fs.trans@exprs[,1]) * 100, digits = 3)
            } else {
                fAI_rem <- 0
                Time.flowAI <- 0
            }
            if (skip_flowClean == F){
                fClean_rem <- round((length(fs.trans@exprs[,1])-length(f.clean@exprs[,1]))/length(fs.trans@exprs[,1]) * 100, digits = 3)
            } else {
                fClean_rem <- 0
                Time.flowClean <- 0
            }
        }
    return( t(c(
        paste0(fileNames[x1]),
        round(F1_rand,digits = 3),round(F1_fcut,digits = 3),round(F1_fAI,digits = 3),round(F1_fclean,digits = 3),
        round(recall_rand,digits = 3),round(recall_fcut,digits = 3),round(recall_fAI,digits = 3),round(recall_fclean,digits = 3),
        round(precision_rand,digits = 3),round(precision_fcut,digits = 3),round(precision_fAI,digits = 3),round(precision_fclean,digits = 3),
        Time.flowCut, Time.flowAI, Time.flowClean,
        resTable[16],Manu_rem,Rand_rem,fCut_rem,fAI_rem,fClean_rem))
    )
}

######## save results and calculate numbers for paper ########
header <-  c("Name",
             "F1_rand", "F1_fCut", "F1_fAI", "F1_fClean",
             "recall_rand","recall_fCut", "recall_fAI", "recall_fClean",
             "precision_rand","precision_fCut", "precision_fAI",   "precision_fClean",
             "time_fCut", "time_fAI", "time_fClean",
             "pass_or_flag_fCut", "%rem_Manu", "%rem_rand", "%rem_fCut", "%rem_fAI", "%rem_fClean")

colnames(results) <- header

write.csv(results, file=paste0(getwd(), "/results/55/F1_scores_results_category_two.csv"))

row1 <- c(mean(as.numeric(results[,'F1_rand'])), mean(as.numeric(results[,'F1_fCut'])), mean(as.numeric(results[,'F1_fAI'])), mean(as.numeric(results[,'F1_fClean'])))

row2 <- c(mean(as.numeric(results[,'recall_rand'])), mean(as.numeric(results[,'recall_fCut'])), mean(as.numeric(results[,'recall_fAI'])), mean(as.numeric(results[,'recall_fClean'])))

row3 <- c(mean(as.numeric(results[,'precision_rand'])), mean(as.numeric(results[,'precision_fCut'])), mean(as.numeric(results[,'precision_fAI'])), mean(as.numeric(results[,'precision_fClean'])))

temp1 <- str_split(results[,'time_fCut'], pattern=":")
tmp1 <- sum(sapply(temp1, function(x) {360*as.numeric(x[1]) + 60*as.numeric(x[2]) + as.numeric(x[3])}))/nrow(results)
temp2 <- str_split(results[,'time_fAI'], pattern=":")
tmp2 <- sum(sapply(temp2, function(x) {360*as.numeric(x[1]) + 60*as.numeric(x[2]) + as.numeric(x[3])}))/nrow(results)
temp3 <- str_split(results[,'time_fClean'], pattern=":")
tmp3 <- sum(sapply(temp3, function(x) {360*as.numeric(x[1]) + 60*as.numeric(x[2]) + as.numeric(x[3])}))/nrow(results)
row4 <- c(0, tmp1, tmp2, tmp3)

row5 <- c(mean(as.numeric(results[,'%rem_rand'])), mean(as.numeric(results[,'%rem_fCut'])), mean(as.numeric(results[,'%rem_fAI'])), mean(as.numeric(results[,'%rem_fClean'])))


F1.scores.table <- cbind(row1, row2, row3, row4, row5)
rownames(F1.scores.table) <- c("random", "flowCut", "flowAI", "flowClean")
colnames(F1.scores.table) <- c("F1 scores", "Recall", "Precision", "Run times in seconds", "Percentages removed")

print(F1.scores.table)

write.csv(F1.scores.table, file =paste0(getwd(), "/results/55/F1_scores_table_category_two.csv") )

cat("Total code time is: ",TimeOutput(start1),"\n",sep="")
```
