## Supplementary material for "flowCut — An R package for precise and accurate automated removal of outlier events and flagging of files based on time versus fluorescence analysis": paper_code: flowCut_Cleaning_comparison_bytype.R

```
library("flowCore") # 1.51.7
library("flowDensity") # 1.19.5
library("flowCut") # 0.99.12
library("flowClean") # 1.22.0
library("flowAI") # 1.12.7
library("doMC") # 1.3.6
library("foreach") # 1.4.7
library("Cairo") # 1.5-10
library("FlowRepositoryR") # 1.14.1
require("R.utils") # 2.9.0
library("stringr") # 1.4.0

#setwd("/code/Projects/flowCut_Paper/codes_plus/")
 source("code/flowCut_Cleaning_Comparison_Functions.R")
start1 <- Sys.time()
set.seed(5678) # flowAI

no_cores <- detectCores() - 1
registerDoMC(no_cores)

skip_flowClean <- F
skip_flowAI <- F
skip_Plotting <- T

flowCutFailed <- NULL
flowCleanFailed <- NULL
flowAIFailed <- NULL

flowCutTimeout <- NULL
flowCleanTimeout <- NULL
flowAITimeout <- NULL

fileNames <- dir(paste0(getwd(), "/data/FlowRepFiles"), pattern = ".fcs", recursive = T, full.names = F)
allFiles <- dir(paste0(getwd(), "/data/FlowRepFiles"), pattern = ".fcs", recursive = T, full.names = T)
fileNames <- unlist(lapply(fileNames,function(fN){unlist(strsplit(fN,split="/"))[2]}))

gating_thresholds <- unname(as.matrix(read.csv(paste0(getwd(), "/data/FlowRepFiles/Manual_Gates_byProblemtype.csv"))))
score.table <- NULL

# for(x in 14:length(unique(gating_thresholds[,1]))){
 results <- foreach (x = 1:length(unique(gating_thresholds[,1])), .combine = rbind, .maxcombine = length(fileNames), .multicombine = T) %dopar% {
#results <- foreach (x = c(1:length(fileNames)), .combine = rbind, .maxcombine = length(fileNames), .multicombine = T) %dopar% {
  try.results <- tryCatch({
   fN <- unique(gating_thresholds[,1])[x]
  ind <- which(fileNames == fN)
  print(fN)
  precision_fAI <- 0
  precision_fcut <- 0
  precision_fclean <- 0
  recall_fAI <- 0
  recall_fcut <- 0
  recall_fclean <- 0
  F1_fcut <- 0
  F1_fAI <- 0
  F1_fclean <- 0
  
  manual_gates <- gating_thresholds[which(gating_thresholds[,1] == fN ),]
  manual_gates <- matrix(manual_gates, ncol=4)
  print(paste0("Starting ", fileNames[ind]))
  fz <- read.FCS(allFiles[ind])
  Desc <- fz@parameters@data$desc
  Name <- fz@parameters@data$name
  NumEvents <- nrow(fz)
  f_org <- fz
  scat.chans <- c(grep (colnames(fz),pattern = "FSC*"),grep (colnames(fz),pattern = "SSC*"))
  if(length(scat.chans)==0){
    f.comp <- fz
  }else{
    f.marg <- removeMargNegs(fz, chans=scat.chans, neg=-1, verbose= F)
    f.comp <- f.marg$f; ind.marg.neg <- f.marg$ind.marg.neg; remove(f.marg)
  }
  ######## Compensation ########
  des <- fz@description
  matrix_ind <- grep(pattern = "spill", x=names(des),ignore.case = T)
  if(length(matrix_ind)==0){
    message("no spill over matrix is found, skipping compensation...")
  } else{
    
    if ( det(des[matrix_ind][[1]])==1 ) {
      message("Check the spillover matrix, it's probably an identity matrix!")
    } else {
      f.comp <- compensate(f.comp, des[matrix_ind][[1]])
    }
    
  }
  ######## Transformation ########
  time.loc <- which(tolower(f.comp@parameters@data$name) == "time"); names(time.loc) <- NULL
  channels.ind <- setdiff(1:length(colnames(f.comp)), c(scat.chans, time.loc)) #locate channels other than scatter channels
  
  lgl <- try (estimateLogicle(f.comp, channels= colnames(f.comp)[channels.ind], m=5))
  if(class(lgl)=="try-error"){
    
    if(!file.exists("filecorrupt.txt")){
      write.table(t(c("file not standard :", fileId, fileNames[x1])),col.names = FALSE, file = "results/55/filecorrupt.txt")
    }
    write.table(t(c("file not standard :", fileId, fileNames[x1])), col.names = FALSE, file = "results/55/filecorrupt.txt", append = TRUE)
    {next}
  }
  fs.trans <- transform(f.comp, lgl)
  
  z <- ceiling(sqrt(length(channels.ind)+2))
 
  
  #find clean regions for manual
  toremove <- NULL
  for(l in 1:length(manual_gates[,2])){
    a  <- which(fs.trans@exprs[,time.loc]>as.numeric(manual_gates[l,3]))
    a1 <- which(fs.trans@exprs[,time.loc]<as.numeric(manual_gates[l,4]))
    toremove <- c(toremove, intersect(a, a1))
  }
  
  cleanRegion <- (1:nrow(fs.trans@exprs))[-toremove]
  
  #### flowCut #####
  segment <- 500
  
  res.flowCut <- try( flowCut(f=fs.trans, Segment=segment, FileID = fN, Directory = paste0(getwd(),"/results/55/flowCutRep_Plots/"),
                              Plot ="None",Verbose=FALSE) )
  resTable <- res.flowCut$data
  f.tvsf <- res.flowCut$frame
  ind.removed <- res.flowCut$ind
  if(length(ind.removed > 0)){
    ind.clean.fCut <- (1:nrow(fs.trans@exprs))[-ind.removed]
  }else{
    ind.removed <- NULL
    ind.clean.fCut <- (1:nrow(fs.trans@exprs))
  }
  
  
  ###### flowAI 
  fAI.cleaned_All <- flow_auto_qc(fs.trans, html_report = FALSE, mini_report = FALSE,
                                  remove_from="FS_FM", second_fractionFR = 0.1, fcs_QC = FALSE, output = 2 )
  
  rem.ind_All <- which(fAI.cleaned_All@parameters@data$name == "remove_from_FS_FM")
  to.be.removed.AI_All <- which(fAI.cleaned_All@exprs[,rem.ind_All] >= 9999)
  if(length(to.be.removed.AI_All) > 0){
    ind.clean.fAI <-  (1:nrow(fs.trans@exprs))[-to.be.removed.AI_All]
  }else{
    to.be.removed.AI_All <- NULL
    ind.clean.fAI <- (1:nrow(fs.trans@exprs))
  }
  
  
  ####flowClean#######
  FSC.loc  <- grep("fsc", tolower(fs.trans@parameters@data$name));  names( FSC.loc) <- NULL
  SSC.loc  <- grep("ssc", tolower(fs.trans@parameters@data$name)); names( SSC.loc) <- NULL
  Time.loc <- which(tolower(fs.trans@parameters@data$name) == "time"); names(Time.loc) <- NULL
  CleanChan.loc <- (1:ncol(fs.trans))[-c(FSC.loc, SSC.loc, Time.loc)]
  res.fclean <- withTimeout({
    try.flowClean <- try(f.clean <- clean(fs.trans, vectMarkers=CleanChan.loc, ext="fcs", filePrefixWithDir=paste("temp/Clean",sep="/")))
    if(class(try.flowClean) =="try-error"){
      f.clean <- fs.trans
      to.be.removed.fC <- NULL
      flowCleanError <- "Error"
      flowCleanFailed <- c(flowCleanFailed, paste0(fileId, "_", x1))
      cat("flowClean with the current values does not work. \n")
      if(!file.exists("results/55/flowCleanError.txt")){
        write.table(t(c("flowClean Error File :", fileId)),col.names = FALSE, file = "results/55/flowCleanError.txt")
      }
      write.table(t(c("flowClean Error File :", fileId)),col.names = FALSE, file = "results/55/flowCleanError.txt", append = TRUE)
    } else {
      rg <- rectangleGate(filterId="gvb", list("GoodVsBad"=c(0, 9999)))
      
          idx <- flowCore::filter(f.clean, rg)
          if(length(which(idx@subSet==F))!=0){
            to.be.removed.fC <- which(idx@subSet==F)
            
            exprs(f.clean) <- exprs(f.clean)[-which(idx@subSet==F),]
            
            
          } else {
            f.clean <- fs.trans
            to.be.removed.fC <- NULL
          }
        
      
    
     
    }
  }, timeout=3600);
  if(length(to.be.removed.fC)==0 ){
    ind.clean.fClean <- (1:nrow(fs.trans@exprs))
  }else{
    ind.clean.fClean <-  (1:nrow(fs.trans@exprs))[-to.be.removed.fC]
  }
  
  
  
  #calculate F1 score for cleaned sections.
  precision_clean_fcut <- length(intersect(ind.clean.fCut,cleanRegion))/length(cleanRegion)
  if(is.nan(precision_clean_fcut)){ precision_clean_fcut <- 0 }
  recall_clean_fcut <- length(intersect(ind.clean.fCut,cleanRegion))/length(ind.clean.fCut)
  if(is.nan(recall_clean_fcut)){ recall_clean_fcut <- 0 }
  F1_clean_fcut <- 2*(precision_clean_fcut *recall_clean_fcut)/(precision_clean_fcut+recall_clean_fcut)
  if(is.nan(F1_clean_fcut)){ F1_clean_fcut <- 0}
  
  precision_clean_fAI <- length(intersect(ind.clean.fAI,cleanRegion))/length(cleanRegion)
  if(is.nan(precision_clean_fAI)){ precision_clean_fAI <- 0 }
  recall_clean_fAI <- length(intersect(ind.clean.fAI,cleanRegion))/length(ind.clean.fAI)
  if(is.nan(recall_clean_fAI)){ recall_clean_fAI <- 0 }
  F1_clean_fAI <- 2*(precision_clean_fAI *recall_clean_fAI)/(precision_clean_fAI+recall_clean_fAI)
  if(is.nan(F1_clean_fAI)){ F1_clean_fAI <- 0}
  
  precision_clean_fClean <- length(intersect(ind.clean.fClean,cleanRegion))/length(cleanRegion)
  if(is.nan(precision_clean_fClean)){ precision_clean_fClean <- 0 }
  recall_clean_fClean <- length(intersect(ind.clean.fClean,cleanRegion))/length(ind.clean.fClean)
  if(is.nan(recall_clean_fClean)){ recall_clean_fClean <- 0 }
  F1_clean_fClean <- 2*(precision_clean_fClean *recall_clean_fClean)/(precision_clean_fClean+recall_clean_fClean)
  if(is.nan(F1_clean_fClean)){ F1_clean_fClean <- 0}
  
  
  F1_score <- matrix(0,ncol=10,nrow=length(manual_gates[,2]))
  #F1 calculations for each problematic region
  for(n in 1:length(manual_gates[,2])){
    toremoveRegion <- NULL
    a  <- which(fs.trans@exprs[,time.loc]>as.numeric(manual_gates[n,3]))
    a1 <- which(fs.trans@exprs[,time.loc]<as.numeric(manual_gates[n,4]))
    toremoveRegion <- c(toremoveRegion, intersect(a, a1))
    #flowCut F1 score calculation
    #find index inside the gating windown
    b <- ind.removed[which(fs.trans@exprs[ind.removed,time.loc] > as.numeric(manual_gates[n,3]))]
    b1 <- ind.removed[which(fs.trans@exprs[ind.removed,time.loc] < as.numeric(manual_gates[n,4]))]
    ind.removed.kept <- c(intersect(b,b1))
    if(length(ind.removed)==0){
      ind.removed <- 0
      ind.removed.kept <- 0
    }
    
    precision_fcut <- length(intersect(ind.removed.kept,toremoveRegion))/length(toremoveRegion)
    if(is.nan(precision_fcut)){ precision_fcut <- 0 }
    recall_fcut <- length(intersect(ind.removed.kept,toremoveRegion))/length(ind.removed.kept)
    if(is.nan(recall_fcut)){ recall_fcut <- 0 }
    F1_fcut <- 2*(precision_fcut*recall_fcut)/(precision_fcut+recall_fcut)
    if(is.nan(F1_fcut)){ F1_fcut <- 0}
    
    F1_score[n,1] <- F1_fcut
    F1_score[n,2] <- precision_fcut
    F1_score[n,3] <- recall_fcut
  
    #flowAI F1 score calculation
    #find index inside the gating windown
    b2 <- to.be.removed.AI_All[which(fs.trans@exprs[to.be.removed.AI_All,time.loc] > as.numeric(manual_gates[n,3]))]
    b3 <-  to.be.removed.AI_All[which(fs.trans@exprs[to.be.removed.AI_All,time.loc] < as.numeric(manual_gates[n,4]))]
    to.be.removed.AI_All.kept <- c(intersect(b2,b3))
    
    fAI.cleaned2_All <- fAI.cleaned_All
    if( length(to.be.removed.AI_All.kept) >= 1 ){
      fAI.cleaned2_All@exprs <- fAI.cleaned_All@exprs[-to.be.removed.AI_All.kept,]
      precision_fAI <- length(intersect(to.be.removed.AI_All.kept,toremoveRegion))/length(toremoveRegion)
      if(is.nan(precision_fAI)){ precision_fAI <- 0 }
      recall_fAI <- length(intersect(to.be.removed.AI_All.kept,toremoveRegion))/length(to.be.removed.AI_All.kept)
      if(is.nan(recall_fAI)){ recall_fAI <- 0 }
      F1_fAI <- 2*(precision_fAI*recall_fAI)/(precision_fAI+recall_fAI)
      if(is.nan(F1_fAI)){ F1_fAI <- 0 }
    }
    
    
    
    F1_score[n,4] <- F1_fAI
    F1_score[n,5] <- precision_fAI
    F1_score[n,6] <- recall_fAI
    
    #flowClean F1 calculation
    b4 <- which(fs.trans@exprs[to.be.removed.fC,time.loc] > as.numeric(manual_gates[n,3]))
    b5 <- which(fs.trans@exprs[to.be.removed.fC,time.loc] < as.numeric(manual_gates[n,4]))
    to.be.removed.fC.kept <- c(intersect(b4,b5))
    precision_fclean <- length(intersect(to.be.removed.fC.kept,toremoveRegion))/length(toremoveRegion)
    if(is.nan(precision_fclean)){ precision_fclean <- 0 }
    recall_fclean <- length(intersect(to.be.removed.fC.kept,toremoveRegion))/length(to.be.removed.fC.kept)
    if(is.nan(recall_fclean)){ recall_fclean <- 0 }
    F1_fclean <- 2*(precision_fclean*recall_fclean)/(precision_fclean+recall_fclean)
    if(is.nan(F1_fclean)){ F1_fclean <- 0 }
    F1_score[n,7] <- F1_fclean 
    F1_score[n,8] <- precision_fclean
    F1_score[n,9] <- recall_fclean
    
   
    F1_score[n,10] <- length(toremoveRegion)/nrow(fs.trans@exprs) #proportions
    
    
  }
  
  temp <- cbind(manual_gates[,1],manual_gates[,2],F1_score,
                paste0(resTable[1],resTable[3],resTable[5],resTable[7]))
  
  F1_clean_score <- c(unique(manual_gates[,1]),"clean",
                      F1_clean_fcut,precision_clean_fcut,recall_clean_fcut,
                      F1_clean_fAI,precision_clean_fAI,recall_clean_fAI,
                      F1_clean_fClean,precision_clean_fClean,recall_clean_fClean,
                      length(cleanRegion)/nrow(fs.trans@exprs),
                      paste0(resTable[1],resTable[3],resTable[5],resTable[7])) 
  
  score.table <- rbind(temp,unname(F1_clean_score))
  },error = function(ex) {
    file.name <- paste0(fN)   
    return(c(file.name, ex))
  })
  return(try.results)
} 

#The file has been flagged. The largest continuous jump was larger than 10% of the range of the 2-98 percentile of the full data.
#The file has been flagged. The max ranged means differ more than 15% of the range of the 2-98 percentile of the full data.
header <-  c("Name","Type",
             "F1_fCut","precision_fCut", "recall_fCut",  
             "F1_fAI", "precision_fAI","recall_fAI",
             "F1_fClean","precision_fClean", "recall_fClean",
             "%rem_Manu","pass_or_flag_fCut")
colnames(results) <- header

write.csv(results, file=paste0("~/results/55/F1_scores_results_byProblemType.csv"))
#write.csv(results, file=paste0("~/results/55/F1_scores_results_byProblemType_extra.csv"), append =TRUE)
```
