## Supplementary material for "flowCut — An R package for precise and accurate automated removal of outlier events and flagging of files based on time versus fluorescence analysis": paper_code: flowCut_Cleaning_Comparison_Functions.R

```
############################################################################################
# removes all cells that exist on the edges of FSC or SSC and the negative events
# Written by Mernoush Malek (BC Cancer)
removeMargNegs <- function(f,chans, sens=1, neg=500, verbose = T)
{
    neg <- rep(neg, length(chans))
    data <- exprs(f)
    margins <- c()

    # removes max
    margins.total <- NULL
    for(chan in chans)
    {
        if (is.character(chan))
            chan <- which(colnames(f)==chan)
        stain.max <- max(data[,chan])
        margins <- which ( data[, chan] >= stain.max*sens)
        margins.total <- c(margins.total, margins)
        if(verbose == T){print(paste(length(margins), "margin events in",colnames(f)[chan], "will be removed.",sep =" "))}
    }

    # removes negatives
    negs.total <- NULL
    for(i in 1:length(chans))
    {
        if (neg[i]<500)
        {
            negs <- which ( data[, chans[i]] < neg[i])
            negs.total <- c(negs.total, negs)
            if(verbose == T){print(paste(length(negs), "negative events in",colnames(f)[chans[i]], "will be removed.",sep =" "))}
        }
    }

    ind.marg.neg <- sort(union(margins.total, negs.total))

    if (length(ind.marg.neg)!=0)
        data <- data[ -ind.marg.neg, ]

    exprs(f) <- data

    return(list(f=f, ind.marg.neg = ind.marg.neg))
}

#################################################################################
# Prints out the time since start_time. Used for optimizing code.
TimeOutput <- function(start_time) {
    start_time <- as.POSIXct(start_time)
    dt <- difftime(Sys.time(), start_time, units="secs")
    # Since you only want the H:M:S, we can ignore the date...
    # but you have to be careful about time-zone issues
    format(.POSIXct(dt,tz="GMT"), "%H:%M:%S")
}
TimeOutput(Sys.Date())
```
