## Supplementary material for "flowCut — An R package for precise and accurate automated removal of outlier events and flagging of files based on time versus fluorescence analysis": paper_code: helperfunc.R

```
getGlobalFrame <- function(fs, sample.length=NA, r = 2){
  if (is(fs, 'flowFrame')){
    return (frame)
  }
  n <- length(fs)
  sample.n <- ifelse(is.na(sample.length),n,sample.length)
  global.frame <- fsApply(fs[sample(n, sample.n)], 
                          function(frame){
                            m <- nrow(frame)
                            exprs(frame )<- exprs(frame)[sample(m, min(m, round(m*r/(sample.n*2)))),]
                            return (frame)
                          })
  global.frame <- as(global.frame, 'flowFrame')
  return (global.frame)
}
averageGates <- function(vec, med =T, global.var= NA,sd.coeff = 2){
  # Args:
  #   vec: a numeric vector of values
  #   med: to use median 
  #   global.var: a number used instead of median
  # Value:
  #   vec with outlying values replaced by the median
  vec <- as.vector(vec)
  m <- ifelse(med,median(vec,na.rm=T),global.var)
  vec[is.na(vec)] <- m
  N <- length(vec)
  c4 <- sqrt(2/(N-1)) * gamma(N/2)/gamma((N-1)/2)
  sdev <- sd(vec,na.rm=T)/c4
  outliers <- which(abs(vec - m)/sdev > sd.coeff)
  if (any(outliers)){
    cat("Changing outliers", outliers, ".\n")
    vec[outliers] <- median(vec[-outliers])
  }
  return (vec)
}
which.index <- function(frame){
  
  return(as(which(!is.na(exprs(frame[,1]))),Class="list"))
  
}
save.index <- function(ind.list,dir)
{
  lapply(1:length(ind.list), function(x) {indices <- lapply(ind.list[[x]], unlist)
                                          save(indices,file= paste(dir, "/Cellindex-", names(ind.list)[x], ".RData",sep=""))
                                          return(cat( "saving", names(ind.list)[x] ,"indices", "\n"))
                                          })
}
removeMargins<- function(f,chans,Plot=F,res.dir=NA)
{
  data <- exprs(f)
  margins <- c()
  for(chan in chans)
  {
    stain.max <-max(exprs(f)[,chan])
    margins <- append(margins,which ( data[, chan] >= stain.max))
   
  }
  cat(length(unique(margins)), "Cells removed as margins in ",chans, "\n")
  new.data <- data[ -unique(margins), ]
  exprs(f) <- new.data
  if(Plot & length(chans)==2)
  {
    if(!is.na(res.dir))
    {
      png(paste(res.dir,identifier(f),"_Margins.png",sep=""),width=900,height=900,pointsize=23)
      }else{
        x11(width=8,height=8)
      }
    inds <- match(chans, colnames(f))
    x.range <- c(max(0,pData(f@parameters)$minRange[inds[1]]),pData(f@parameters)$maxRange[inds[1]])
    y.range <- c(max(0,pData(f@parameters)$minRange[inds[2]]),pData(f@parameters)$maxRange[inds[2]])
   plot(data[,chans],pch=".", xlab=chans[1],ylab=chans[2],col=1,xlim=x.range,ylim=y.range, main=paste(length(unique(margins)),"margins"))
     points(data[margins, chans],pch=".",col=2)
    dev.off()
  }
  
  return(f)
}
Make.FCS<- function(markers, data)
{
  pd <- c()  # 'params' phenoData
  des <- list()  # 'description' list
  
  des[["$DATATYPE"]] <- "F"
  for (c in 1:ncol(data)) {
    c_name <- colnames(data)[c]
    c_marker<-markers[c]
    c_min <- floor(min(data[,c]))
    c_max <- ceiling(max(data[,c]))
    c_rng <- c_max - c_min + 1
    
    pl <- matrix(c(c_name, c_marker, c_rng, c_min, c_max),nrow=1)
    colnames(pl) <- c("name", "desc", "range", "minRange", "maxRange")
    rownames(pl) <- paste("$P",c,sep="") 
    pd <- rbind(pd, pl)
    
    des[[paste("$P",c,"B",sep="")]] <- "32";      # Number of bits
    des[[paste("$P",c,"R",sep="")]] <- toString(c_rng); # Range
    des[[paste("$P",c,"E",sep="")]] <- "0,0";      # Exponent
    des[[paste("$P",c,"N",sep="")]] <- c_name;	    # Name
    des[[paste("$P",c,"S",sep="")]] <- c_marker;	    # Desc	
  }
  frame<-flowFrame(data, as(data.frame(pd), "AnnotatedDataFrame"), description=des)
  return(frame)
}
estimate.logicle<-function(fs,talk= TRUE,return.set=TRUE,med=TRUE, trans.chans=NULL,estimate=T)
  {
  f<-fs[[1]]
  if (is.null(trans.chans))
  {
   log.channels <- paste("P",1:ncol(f),"DISPLAY",sep="")
   trans.chans <- which(f@description[log.channels]=="LOG")
   if(any(is.null(trans.chans)) | any(is.na(trans.chans)) | (length(trans.chans)==0))
      {
    return(cat("You have to find the channels manually, there's no information on FCS file","\n"))
   }
  }
  if( talk)
    cat("Channels to be transformed ", trans.chans, "\n")
  if(estimate)
  {
  if (med==TRUE){
    lgl <-estimateMedianLogicle(flow_set=fs,channels=colnames(f)[trans.chans])
   
  }else{
     lgl<-estimateLogicle(getGlobalFrame(fs),channels=colnames(f)[trans.chans])
  }
  if(return.set) 
  { 
 
     return(fsApply(fs,function(x) return(transform(x,lgl))))
  }else
    return(lgl)
}else{
  library(Logicle)
  trans.fs <-fsApply(fs,function(frame) {
    lg<-Logicle::create(T=262144, W=0.5)
    x<-exprs(frame)[,trans.chans]
    y<-Logicle::scale(lg, x)
    exprs(frame)[,trans.chans]<-y
   # detach(package:Logicle, unload=TRUE)
    return (frame)
  })
  }
}

QA.process <- function(fs,dir, call.pdf=FALSE){
  
  cell.num <-qaProcess.cellnumber(fs, grouping=NULL, outdir=dir, cFactor=2,
                                  absolute.value=NULL, two.sided = FALSE,
                                  name="cell number", sum.dimensions=NULL,
                                  pdf=call.pdf)
  margin <- qaProcess.marginevents(fs, channels=colnames(fs)[c(1,4,7:24)],side ="both", grouping=NULL, 
                                   outdir=dir, cFactor=2, absolute.value=NULL, name="margin events",
                                   sum.dimensions=NULL, det.dimensions=c(3,1), pdf=call.pdf)
  writeQAReport(fs, processes=list(cell.num,margin), globalProcess=NULL, 
                outdir = paste(dir,"/qaReport",sep=""), grouping = NULL,pagebreaks=TRUE, pdf=call.pdf)
  
  
}

rotate.data <- function(data, chans=NULL, theta=NULL)
{
  if (class(data)== "flowFrame" & !is.null(chans))
  { 
    data.new <- exprs(data)[,chans]
    if (is.null(theta))
    {
      reg.slope <- atan(lm(data.new[,1] ~ data.new[,2])$coefficients[2])
      theta <- pi/2 - reg.slope
    }
  data.new <- data.new %*% matrix(c(cos(theta),-sin(theta),sin(theta),cos(theta)),2,2,byrow=T)
  exprs(data)[,chans] <- data.new
  }else{
    data <- data %*% matrix(c(cos(theta),-sin(theta),sin(theta),cos(theta)),2,2,byrow=T)
  }
  return(list(data=data,theta=theta))
}
plot.layout <- function(x)
{
  if(round(sqrt(x))^2 >= x)
    n<-m <- round(sqrt(x))
  else{
    n<- round(sqrt(x))
    m <- round(sqrt(x))+1
  }
  return (c(m,n))
}
#' Estimates a common logicle transformation for a flowSet.
#'
#' Of the negative values for each channel specified, the median of the specified
#' quantiles are used.
#'
#' @param flow_set object of class 'flowSet'
#' @param channels character vector of channels to transform
#' @param m TODO -- default value from .lgclTrans
#' @param q quantile
#' @return TODO
estimateMedianLogicle <- function(flow_set, channels, m = 4.5, q = 0.05) {
  if (!is(flow_set, "flowSet")) {
    stop("flow_set has to be an object of class 'flowSet'")
  }
  if (missing(channels)) {
    stop("Please specify the channels to be logicle transformed")
  }
  indx <- channels %in% unname(colnames(exprs(flow_set[[1]])))
  if (!all(indx)) {
    stop(paste("Channels", channels[!indx], "were not found in flow_set "))
  }
  
  neg_marker_quantiles <- fsApply(flow_set, function(sample) {
    apply(exprs(sample), 2, function(markers) {
      quantile(markers[markers < 0], probs = q)
    })
  })
  # Replaces 'r' in flowCore:::.lgclTrans
  neg_marker_quantiles <- apply(neg_marker_quantiles, 2,
                                median, na.rm = TRUE)[channels]
  
  # In the case that no negative markers are present, we set this quantile to the
  # default value of 1/2.
  neg_marker_quantiles <- replace(neg_marker_quantiles,
                                  is.na(neg_marker_quantiles), 0.5)
  
  # Replaces 't' in flowCore:::.lgclTrans
  max_range <- do.call(rbind, lapply(fsApply(flow_set, range), function(x) {
    x[2, channels]
  }))
  max_range <- apply(max_range, 2, max)
  
  # Replaces 'w' in flowCore:::.lgclTrans
  w <- (m - log10(max_range / abs(neg_marker_quantiles))) / 2
  
  transformation <- lapply(channels, function(channel) {
    transId <- paste(channel, "medianLogicleTransform", sep = "_")
    
    logicleTransform(transformationId = transId, w = w[channel],
                     t = max_range[channel], m = m, a = 0)
  })
  
  transformList(channels, transformation,
                transformationId = "medianLogicleTransform")
}
# getPeaks <- function(frame,chans,tinypeak.removal=tinypeak.removal){
#   
#   ##=====================================================================================================================
#   ## Finds the peaks in the given density
#   ## Args:
#   ##   dens: density of the channel whose peaks are going to be found. It is a variable of class 'density'.
#   ##   w: the length of the window where the function searches for the peaks. If all peaks required, use the default w=1.
#   ## Value:
#   ##   peaks in the density of the provided channel
#   ##---------------------------------------------------------------------------------------------------------------------
#   data <- exprs(frame)[,chans]
#   dens <- density(data[which(!is.na(data))])
#   dens <- smooth.spline(dens$x, dens$y, spar=0.4)
#   dens$y[which(dens$y<0)] <- 0
#   d <- dens$y
#   w <- 1
#   peaks <- c()
#   peaks.ind <- c()
#   for(i in 1:(length(d)-w)){
#     if(d[i+w] > d[(i+w+1):(i+2*w)] && d[i+w] > d[i:(i+w-1)] && d[i+w] > tinypeak.removal*max(d)){ # also removes tiny artificial peaks less than ~%4 of the max peak
#       peaks <- c(peaks, dens$x[i+w])
#       peaks.ind <- c(peaks.ind, i+w)
#     }
#   }
#   return(list(Peaks=peaks, Dens=dens,Ind=peaks.ind))
# }
getIntersect <- function(dens, p1, p2){
  
  ##=================================================================
  ## Returns the min intersection between two peaks
  ## Args:
  ##   dens: density of the channel whose peaks are going to be found
  ##   p1: firs peak
  ##   p2: second peak
  ## Value:
  ##   the min intersection between two peaks
  ##-----------------------------------------------------------------
  
  if(missing(p2) | is.na(p2))
    p2 <- min(dens$x)
  fun <- splinefun(dens)
  index <- intersect(which(dens$x < max(c(p1,p2))), which(dens$x > min(c(p1,p2))))
  min.intsct <- dens$x[index][which.min(fun(dens$x[index]))]
  #if(length(min.intsct==0) | is.infinite(min.intsct))
  #  min.intsct <- min(dens$x)
  return(min.intsct)
}
```
