## Supplementary material for "flowCut — An R package for precise and accurate automated removal of outlier events and flagging of files based on time versus fluorescence analysis": paper_code: IMPC_Functions.R

```
############################################################################################
# removes all cells that exist on the edges of FSC or SSC and the negative events
removeMargNegs<- function(f,chans, sens=1, neg=500, verbose = T)
{
    neg <- rep(neg, length(chans))
    data <- exprs(f)
    margins <- c()

    # removes max
    margins.total <- NULL
    for(chan in chans)
    {
        if (is.character(chan))
            chan <- which(colnames(f)==chan)
        stain.max <- max(data[,chan])
        margins <- which ( data[, chan] >= stain.max*sens)
        margins.total <- c(margins.total, margins)
        if(verbose == T){print(paste(length(margins), "margin events in",colnames(f)[chan], "will be removed.",sep =" "))}
    }

    # removes negatives
    negs.total <- NULL
    for(i in 1:length(chans))
    {
        if (neg[i]<500)
        {
            negs <- which ( data[, chans[i]] < neg[i])
            negs.total <- c(negs.total, negs)
            if(verbose == T){print(paste(length(negs), "negative events in",colnames(f)[chans[i]], "will be removed.",sep =" "))}
        }
    }

    ind.marg.neg <- sort(union(margins.total, negs.total))

    if (length(ind.marg.neg)!=0)
        data <- data[ -ind.marg.neg, ]

    # print(data)
    # print(ind.marg.neg)
    # print(dim(data))
    # print(length(ind.marg.neg))

    exprs(f) <- data
    # f@exprs <- data

    return(list(f=f, ind.marg.neg = ind.marg.neg))
}
#################################################################################
# change the date format
changeDate <- function(dates) {
    dates2 <- NULL
    for ( date in  dates){
        date <- gsub( "-14","-2014", date)
        date <- gsub( "-15","-2015", date)
        date <- gsub( "-16","-2016", date)
        date <- gsub("Jan", "01", date)
        date <- gsub("Feb", "02", date)
        date <- gsub("Mar", "03", date)
        date <- gsub("Apr", "04", date)
        date <- gsub("May", "05", date)
        date <- gsub("Jun", "06", date)
        date <- gsub("Jul", "07", date)
        date <- gsub("Aug", "08", date)
        date <- gsub("Sep", "09", date)
        date <- gsub("Oct", "10", date)
        date <- gsub("Nov", "11", date)
        date <- gsub("Dec", "12", date)

        dateComponents <- strsplit(date, '-', fixed=TRUE)[[1]]
        date <- paste0(dateComponents[3], dateComponents[2], dateComponents[1])
        dates2 <- c(dates2, date)
    }
    return(dates2)
}
#################################################################################
# Prints out the time since start_time. Used for optimizing code.
TimeOutput <- function(start_time) {
    start_time <- as.POSIXct(start_time)
    dt <- difftime(Sys.time(), start_time, units="secs")
    # Since you only want the H:M:S, we can ignore the date...
    # but you have to be careful about time-zone issues
    format(.POSIXct(dt,tz="GMT"), "%H:%M:%S")
}
TimeOutput(Sys.Date())

#######################################################################################
## This part was taken from the helperfunc.R script
##Finds markers in the FCS file
Find.markers <- function(frame,marker.list)
{
    #Parameters:
    #*frame: a flowFrame in the flowSet
    #**marker.list: A vector of characters
    #Output:
    #*channels.ind: a vector of channels numbers in the frame  corresponding to marker.list
    channels.ind <- unlist(lapply(marker.list, function(x) {
        ind <- grep(x, frame@parameters@data[,2], ignore.case=T)
        ind_store <- ind
        if(length(ind)==0){
            warning(paste (x, "not found, check markers!"))
            return(NA)
        } else {
            if(length(ind)>1) {
                cnt <- 0
                repeat{
                    cnt <- cnt + 1
                    fs.markers<-unlist(lapply(frame@parameters@data[,2], function(x) unlist(strsplit(x," "))[cnt]))
                    ind<-match(x,fs.markers)
                    if (is.na(ind))
                    {
                        fs.markers<-unlist(lapply(frame@parameters@data[,2], function(x) unlist(strsplit(x,"-"))[cnt]))
                        ind<-match(x,fs.markers)
                        if(!is.na(ind))
                            break;
                    } else {
                        break;
                    }
                    if(cnt >= 10) {

                        if (length(ind_store) >= 2){
                            ind <- ind_store[1]
                            warning(paste (x, "found more than one, choosing first. Check markers!"))
                        } else {
                            warning(paste (x, "not found, check markers!"))
                        }
                        break;
                    }
                }
            }
        }
        return(ind)
    }))
    names(channels.ind)<-marker.list
    #Removing NAs in the channel vector, as not all centres have all of these markers
    #Note that most centres should have Live/CD4/CD8/CD44/CD62/CD5/CD161/CD25 in their channels
    ind <- which (is.na(channels.ind))
    if (length(ind)!=0)
        channels.ind <- channels.ind[-ind]
    return(channels.ind)
}
```
